# Genomic evidence of an early evolutionary divergence event in wild *Saccharomyces cerevisiae*

**DOI:** 10.1101/2020.06.03.131607

**Authors:** Devin P Bendixsen, Noah Gettle, Ciaran Gilchrist, Zebin Zhang, Rike Stelkens

## Abstract

Comparative genome analyses have suggested East Asia to be the cradle of the domesticated microbe Brewer’s yeast (*Saccharomyces cerevisiae*), used in the food and biotechnology industry worldwide. Here, we provide seven new, high quality long read genomes of non-domesticated yeast strains isolated from primeval forests and other natural environments in China and Taiwan. In a comprehensive analysis of our new genome assemblies, along with other long read *Saccharomycetes* genomes available, we show that the newly sequenced East Asian strains are among the closest living relatives of the ancestors of the global diversity of Brewer’s yeast, confirming predictions made from short read genomic data. Three of these strains (termed the East Asian Clade IX Complex here) share a recent ancestry and evolutionary history suggesting an early divergence from other *S. cerevisiae* strains before the larger radiation of the species, and prior to its domestication. Our genomic analyses reveal that the wild East Asian strains contain elevated levels of structural variations. The new genomic resources provided here contribute to our understanding of the natural diversity of *S. cerevisiae*, expand the intraspecific genetic variation found in this this heavily domesticated microbe, and provide a foundation for understanding its origin and global colonization history.

**Significance statement:** Brewer’s yeast (*Saccharomyces cerevisiae*) is a domesticated microbe and research model organism with a global distribution, and suspected origin in East Asia. So far only limited genomic resources are available from non-domesticated lineages. This study provides seven new, high quality long read genomes of strains isolated from primeval forests and other natural environments in China and Taiwan. Comparative genomics reveal elevated levels of structural variation in this group, and early phylogenetic branching prior to the global radiation of the species. These new genomic resources expand our understanding of the evolutionary history of Brewer’s yeast, and illustrate what the ancestors of this highly successful microbe may have looked like.

## Introduction

The history of Brewer’s yeast, *Saccharomyces cerevisiae*, is deeply interwoven with that of humanity, having played significant roles in cultural, technological, and societal development for at least 9000 years (McGovern, et al. 2004). While over a hundred years of *S. cerevisiae* research has provided important insights into eukaryotic genomics, evolution and cell physiology, much of its ‘wild’ ecology as well as its deep human and pre-human evolutionary history have, until recently, largely remained a mystery. Recent broadscale genomic surveys of *S. cerevisiae* and its close relatives, however, are beginning to shed light on important aspects of its population genetic structure, intra- and inter-specific hybridization events, and their interplay in yeast domestication (Duan, et al. 2018; Peter, et al. 2018; Scannell, et al. 2011; Wang, et al. 2012).

One of the key results from these broadscale genomic surveys has been increasing evidence for a singular and central radiation event of *S. cerevisiae* from Far East Asia (Wang, et al. 2012; Peter, et al. 2018; Duan, et al. 2018). These studies have independently revealed that strains of wild yeast collected in parts of China and Taiwan contain much higher genomic diversity and show greater levels of divergence than all other strains of *S. cerevisiae*. The vast majority of these *S. cerevisiae* genomes, however, have been analyzed using short-read sequencing, resulting in a focus on single nucleotide variants (SNVs). Larger structural variations (SVs), such as inversions, deletions and gene duplications, in addition to repetitive regions such as transposable elements (TE) and telomeres, have gone largely unresolved (Goodwin, et al. 2016). In addition to playing significant roles in yeast adaptation (Payen, et al. 2014; Steenwyk and Rokas 2018; Zhang, et al. 2020), these large structural features can provide increased phylogenetic resolution and key insights about lineage interactions and potential reproductive isolation. SVs have shown to be vital for evolutionary adaptation in many other taxa, supporting the role of inversions in adaptation and speciation, and in the evolution of disease (Merker, et al. 2018; Wellenreuther, et al. 2019).

In this study, we generated high quality assemblies of seven of the highly divergent wild East Asian strains and one common laboratory strain (Y55) using both short-reads and PacBio long-reads to better understand the relationships of these strains to the global diversity of *S. cerevisiae*. Analyzing our assemblies in the context of publicly available long-read genomes, we generated a new phylogeny that confirms the place of these East Asian strains at the base of *S. cerevisiae*, and provide further evidence for an out-of-China colonization history of this species. Moreover, we were able to group our sequenced strains belonging to the previously identified CHN IX clade with a Taiwanese strain, both shown in separate studies to be divergent from the rest of *S. cerevisiae*. We show that this combined clade likely has deep roots in mainland China and has had little gene flow with other *S. cerevisiae* strains.

## Results

### Genome sequencing and assembly

We used whole-genome long-read PacBio sequencing to assemble the genomes of seven highly divergent and one common lab strain of *S. cerevisiae* (**Fig. S1**, average per base genomic coverage = 91.3; average median read length = 3028bp). Initial nuclear and mitochondrial assemblies were highly complete (median # of contigs = 25; median N50 = 821424.5). Final nuclear and mitochondrial assemblies were further resolved to single contigs for each chromosome (**Table S1**, median N50 = 907965.5). Final genome sizes ranged from 11.65 to 11.92 Mbp. Assessment of the completeness of the genome assembly and annotation using BUSCO found that all genomes had similarly high BUSCO scores (C > 96.5%, **Table S2**).

### Phylogenomics

Our newly constructed consensus species tree, placed six of the newly assembled East Asian strains in a basal position within the *S. cerevisiae* radiation (Fig. 1). Three of these strains, EM14S01-3B (Taiwanese) (Peter, et al. 2018), XXYS1.4 and JXXY16.1 (CHN IX) (Duan, et al. 2018), hereon referred to as the East Asian Clade IX Complex, show early divergence from all other *S. cerevisiae* strains. Despite the largely basal placement of our assembled East Asian strains, one strain (BJ4) clustered separately with Y12 and YPS128, strains isolated from Ivory Coast palm wine and Pennsylvanian woodland soil, respectively. The common lab strain, Y55, clustered with two other domesticated strains (DBVPG6044 and SK1) within the West African+ clade. Construction of an Alignment and Assembly-Free (AAF) phylogeny comparing the long-read sequencing data generated in this study and previous short-read data found a high-level of similarity between the two datasets (**Fig. S2-S3**). This analysis also found similar clustering to the consensus species tree, among the East Asian strains and the common lab strain as well as a large amount of divergence of the East Asian Clade IX Complex from the rest of *S. cerevisiae.* However, AAF was unable to resolve the deep early divergence of the East Asian Clade IX Complex from other strains.

**Figure 1:**
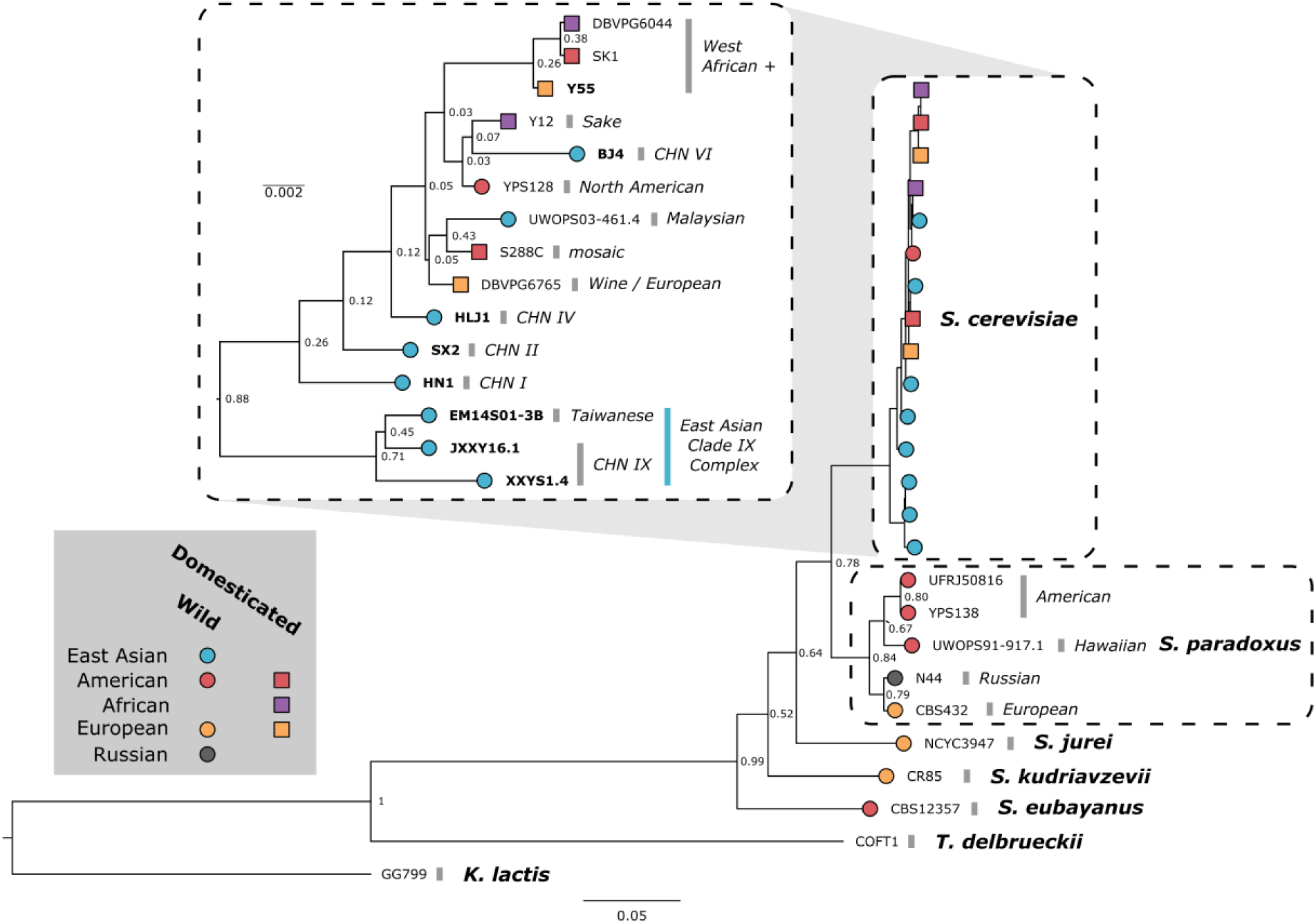
Consensus phylogenetic tree of yeast long-read genomes. The tree was built by orthogroup inference. The support values are the proportion of times that the bipartition is seen in each of the individual species tree estimates. Branch lengths represent the average number of substitutions per site across the sampled gene families. For species with more than a single long-read genome assembly (*S. cerevisiae and S. paradoxus*), species clades are indicated in italics. Strains are colored according to their location of origin and branch tip shape indicates whether it is domesticated (square) or wild (circle). Inset depicts *S. cerevisiae* strains with independent scaling. New long-read genome assemblies presented in this study are indicated in bold.

### Structural variation

A comparison of our eight *S. cerevisiae* genomes and previously assembled *Saccharomyces* sensu stricto genomes to the *S. cerevisiae* reference genome (S288C), revealed a high level of collinearity, particularly at larger scales (Fig. 2, **Fig. S4-S10**). We found exceptions to this strict collinearity only in one strain of *S. paradoxus* (previously reported (Yue, et al. 2017)) and in the East Asian Clade IX Complex. All three member strains show a ~80kb terminal translocation from chromosome XI to chromosome XII (Fig. 2A inset). This structural variant in the East Asian Clade IX Complex was further supported by both long- and short-read analyses of alignment coverage (**Fig. S11-S12**). Additional evidence for this unique translocation comes from high short-read coverage of chromosome XII of XXYS1.4, indicating a likely aneuploidy, which extends across the translocated region of chromosome XI (**Fig. S12**). Other notable rearrangements are a large inversion in chromosome X of BJ4 (**Fig. S5**). The common lab strain, Y55, showed a high level of collinearity with only minor deviations from homology (**Fig. S4**).

**Figure 2:**
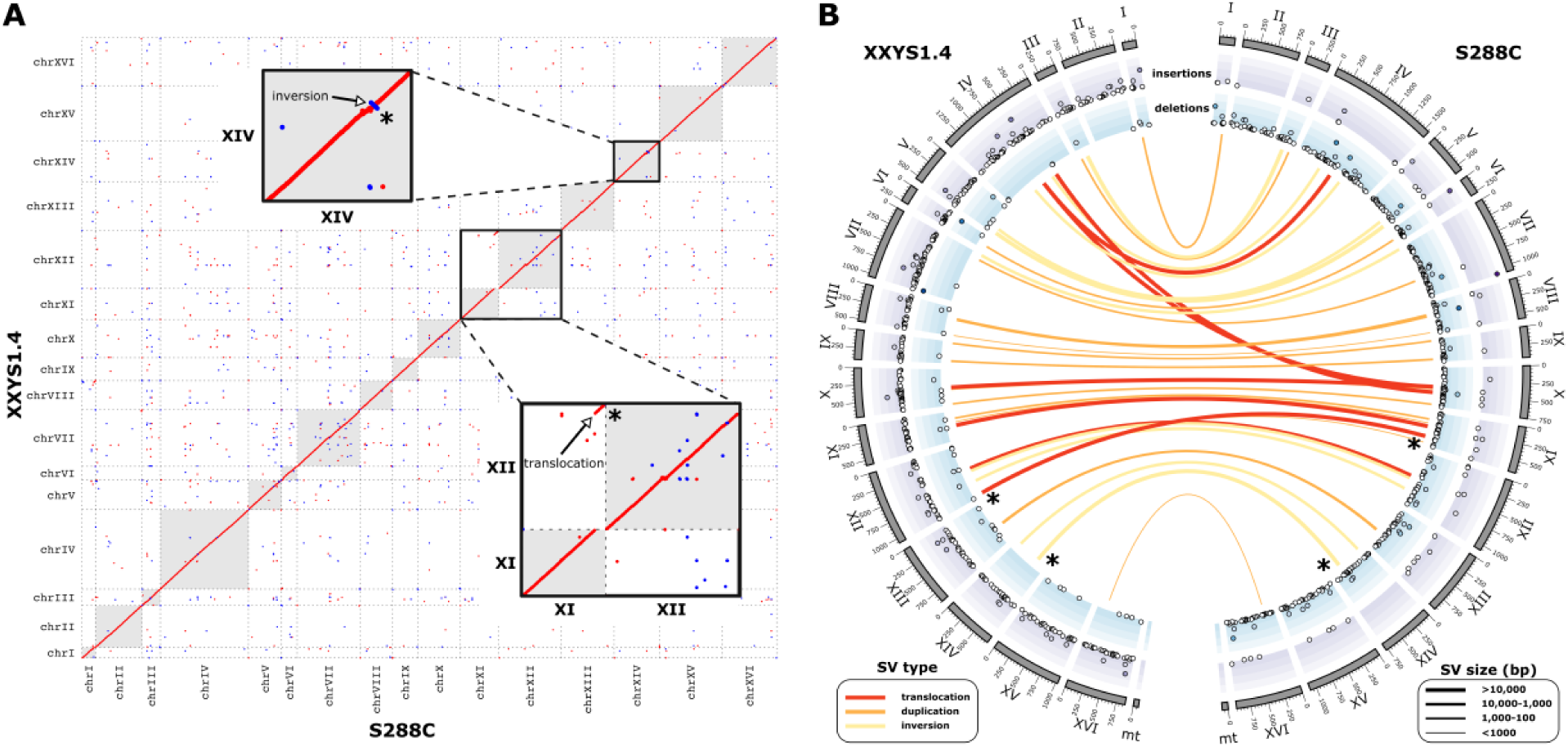
*Saccharomyces cerevisiae* long-read PacBio genome assemblies. **(A)** Genome comparison of the reference strain, S288C, and a member of the East Asian Clade IX Complex: XXYS1.4. Sequence homology within the dot plots is indicated by red dots for forward matches and blue dots for reverse matches. Insets depict examples of deviations from homology: 1) a large translocation between XI and chromosome XII found conserved in JXXY16.1, XXYS1.4 and EM14SO1-3B and 2) a large inversion in chromosome XIV. **(B)** CIRCOS plot showing the detected structural variations between reference strain, S288C and XXYS1.4. Translocations (red), duplications (orange) and inversions (yellow) are depicted as links between the two genomes. The width of the link reflects the relative size of the variation (bp). The translocation and inversion depicted in panel A are highlighted with asterisks. Insertions (blue) and deletions (purple) are depicted in the outer tracks. Deletion and insertion size increase towards the outside. Chromosome size is shown on the outside in 1kb units.

To quantify the extent of smaller structural variations in our genomes, we performed a comprehensive analysis using pairwise comparisons between the 15 *S. cerevisiae* strains with long-read assemblies. We assessed five types of variation: deletions, insertions, duplications, inversions and translocations. This analysis revealed that the wild East Asian strains tend to have higher amounts of total variation (mean=356.5) compared to the other strains (mean=384.7, Fig. 3A). The three Clade IX Complex strains (EM14S01-3B, JXXY16.1, XXYS1.4) were among the highest, and in particular strain XXYS1.4 had a significantly higher mean variant count (525.9, Fig. 4B). In contrast, the common lab strain Y55 had more moderate levels of total variation. The East Asian Clade IX Complex also had larger numbers of deletions and inversions, and fewer insertions and duplications (Fig. 4C, **Fig. S13-S17**). The Malaysian strain UWOPS03-461.4 had significantly larger numbers of translocations compared to all strains. A closer analysis of the distribution of all structural variations identified in this study along chromosomes revealed areas of elevated variation counts, however we found no strong patterns (**Fig. S18**).

**Figure 3:**
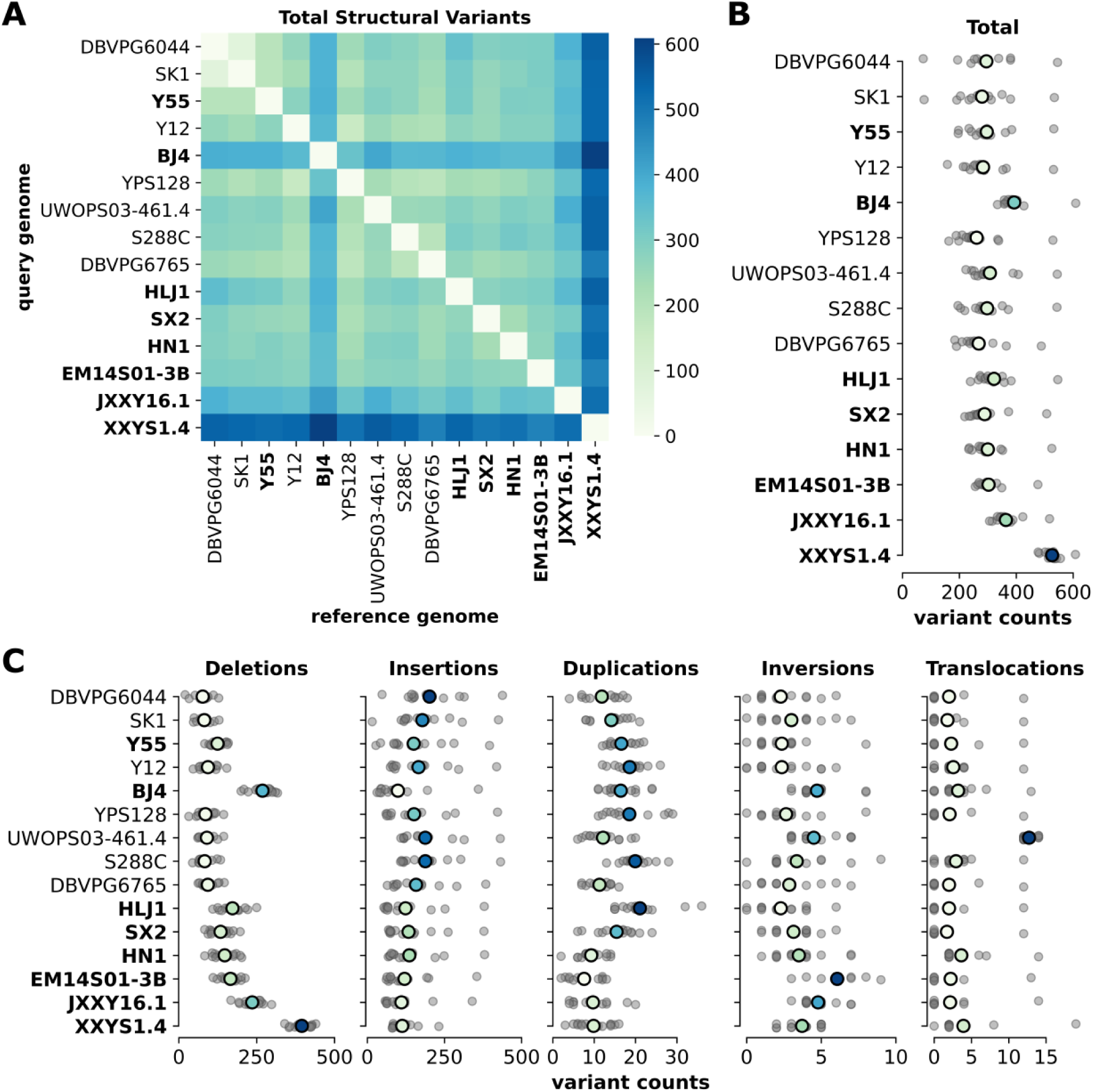
Structural variations within *Saccharomyces cerevisiae*. **(A)** Pairwise comparisons among all ***Saccharomyces cerevisiae*** genome assemblies with the total number of variations. Order of genome assemblies is consistent with the species tree (Fig. 1). New long-read genome assemblies presented in this study are bold. **(B)** The range of total structural variation counts found for each genome serving as reference genome. Grey dots indicate each pairwise genome comparison. Colored dots indicate the mean and are colored on a relative scale. **(C)** The range of structural variation counts for each type of variation. Grey dots indicate each pairwise genome comparison. Colored dots indicate the mean and are colored on a relative scale. Corresponding heatmaps for pairwise comparison are shown in **Fig. S13 to S17**.

**Figure 4:**
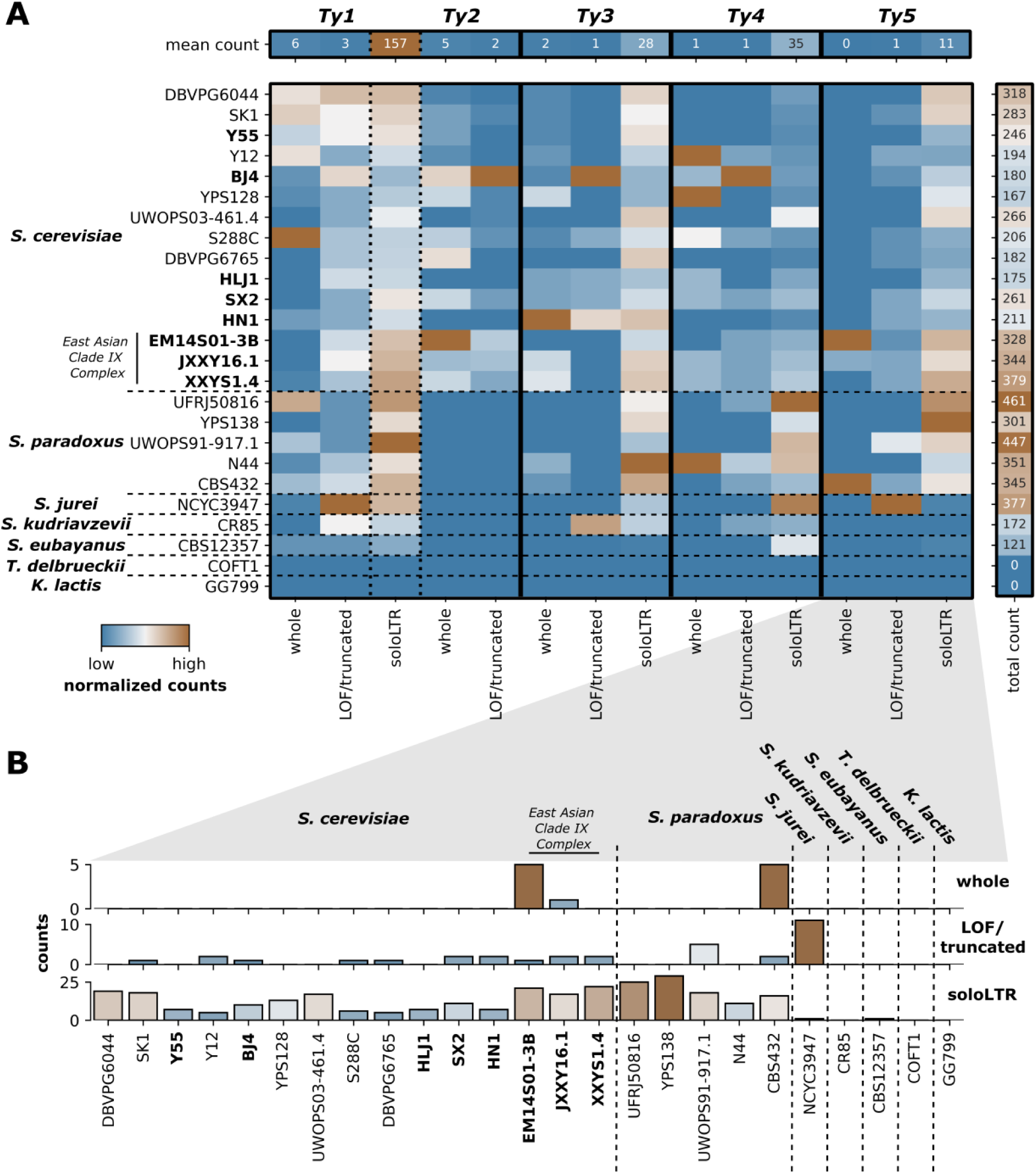
Transposable element composition in *Saccharomyces*. **(A)** Transposable element composition in total count subdivided by *Ty* classification for *Saccharomyces sensu stricto* strains. For visual comparison, each column is normalized (x/x_max_) for that specific element. For raw values see **Fig. S21**. LTR = long terminal repeat components of *Ty* elements without replicative machinery. **(B)** A closer look at *Ty5* elements across *Saccharomyces*.

### Nuclear genome content

In general, our newly assembled long-read genomes were significantly smaller than the currently existing genomes (t = 2.36, df = 10.28, p = 0.039). This difference, however, is largely a result of reduced genome size in members of the East Asian Clade IX Complex (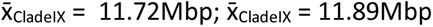; t = 2.93, df = 5.25, p = 0.031). These size differences are due to decreases in genic material both in terms of counts (t = 6.12, df = 10.43, p < 0.001) and cumulative gene length (t = 7.35, df = 5.0, p < 0.001), and a relative reduction in non-coding DNA (t = 4.95, df = 6.74, p = 0.002) (**Fig. S19**). Interestingly, these relative reductions in genic material are correlated with increases in identified intronic material, a pattern that is carried throughout all *S. cerevisiae* strains analyzed here (F = 11.38; r^2^ = 0.43; p = 0.005).

### Transposable Element Composition

Transposable elements (TEs) replicate and deteriorate in a way that gives them an evolutionary history that can be unique with regards to their host genomes and can provide hints about past interactions between distinct lineages. To better understand historical relationships between different strains of *S. cerevisiae*, we annotated and analyzed all classes of known retrotransposon or *Ty* element in this species.

In terms of simple counts, members of the East Asian Clade IX Complex had more *Ty*-associated elements than the rest of the *S. cerevisiae* strains (t = −6.05, df = 6.31, p < 0.001), a result largely based on a disproportionate number of solo long terminal repeats (LTRs) across all classes of *Ty* elements (Fig. 4A, **Fig. S20-S21**). A similar pattern remained when comparing total length of elements (**Fig. S22**). Although *Ty1*/*Ty2* LTRs were the most common *Ty* remnant in all strains, the relative frequency of each class of *Ty* element across *S. cerevisiae* strains does not follow the same pattern reported for the reference strain S288C, where *Ty1* > *Ty2* > *Ty3* > *Ty4* > *Ty5*. Indeed, *Ty1* elements have often been suggested as being the most prolific TE class in *S. cerevisiae*; however, we did not find any putatively functional *Ty1* elements in 6 of the 15 strains we analyze while finding 30 in the reference strain, S288C, representing a clear outlier at the upper end.

As yet, functional *Ty5* elements had only been identified in *S. paradoxus.* “Complete” elements (i.e. elements containing both flanking LTRs and the internal coding region) previously identified in *S. cerevisiae* strains are missing a ^~^2kb portion of the ^~^5kb internal coding region and are found in very low numbers (1-2 per strain). However, the Clade IX Complex strains show a particularly high abundance of *Ty5*-associated elements (Fig. 4B). Further examination revealed six complete *Ty5* elements with fully intact coding regions distributed across two Clade IX Complex strains, EM14S01-3B and JXXY16 (**Fig. S23**). While all “complete” *Ty5* elements that we identified in *S. cerevisiae* outside of the Clade IX Complex are missing the same ^~^2kb region, only 2/10 Clade IX *Ty5* elements (both in JXXY16.1) are missing this region. Additionally, these elements largely do not share homologous bordering regions. In conclusion, the only putatively functional *Ty5* elements in *S. cerevisiae* are in the Clade IX Complex.

### Comparative mitochondrial genomics

Overall, the mitochondrial genomes of the *S. cerevisiae* strains showed high-levels of collinearity (Fig. 5). Of note, however, is the absence of RPM1, a highly conserved ncRNA component of mitochondrial RNase P in two of the Clade IX Complex strains, JXXY16.1 and XXYS1.4. To further confirm the absence of this gene we aligned the reference RPM1 to the unassembled PacBio reads using BLASTn (Zhang, et al. 2000). We found no full-length alignments of RPM1, a 483 bp gene, in either set of reads; rather the highest scoring alignments (e-value>9e-35) were 149 (JXXY16.1) and 239 bp (XXYS1.4). Similarly, we were unable to find or assemble more than a truncated version of the mitochondrial 21s rRNA in JXXY16.1 None of the strains we sequenced were found to be respiratory incompetents or ρ–.

**Figure 5:**
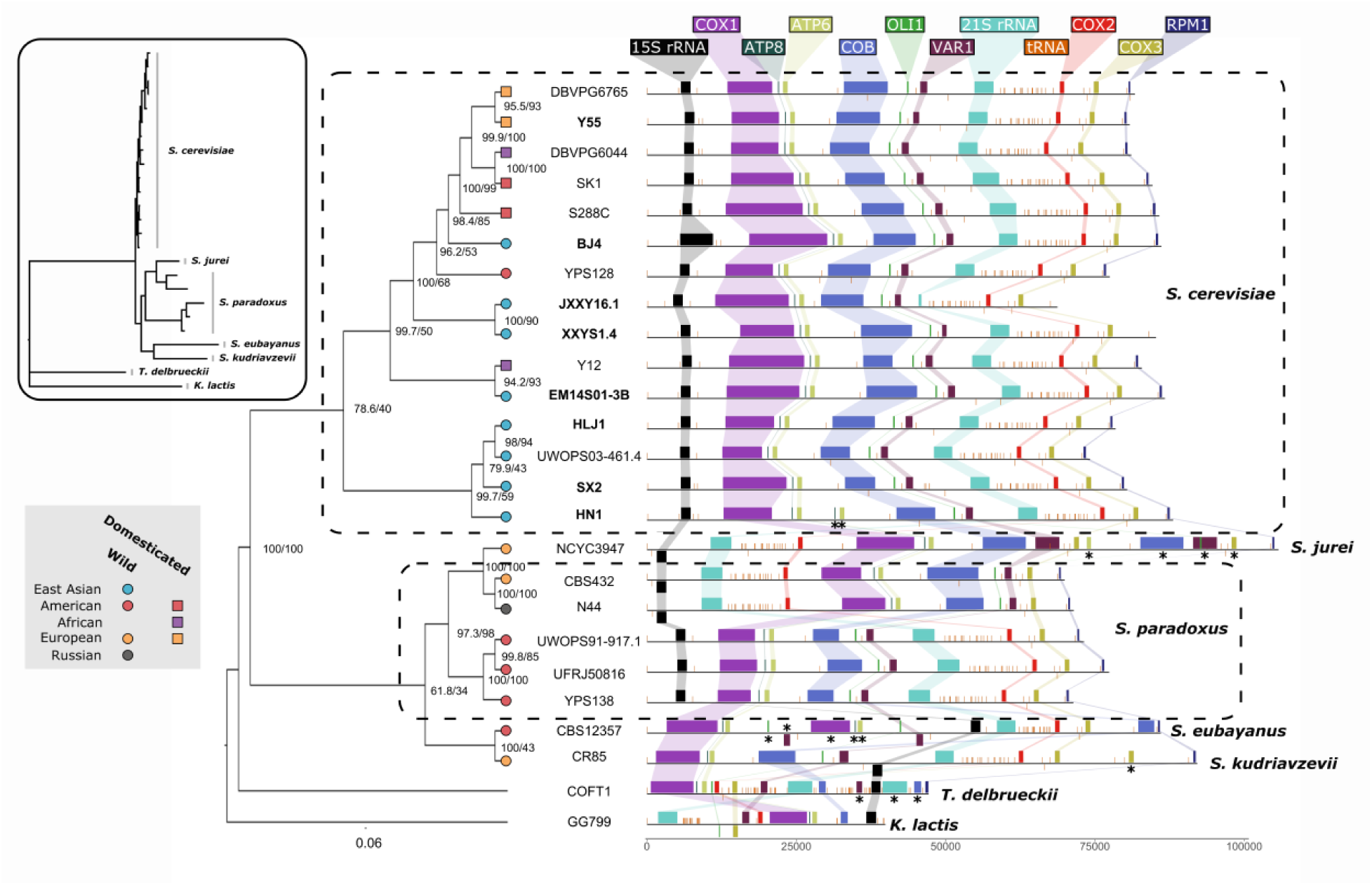
Mitochondrial phylogenetics and genomic arrangements. Phylogenetic tree based on mitochondrial genomic content. Internal branches are labeled with bootstrap support. *Saccharomyces* strains are colored according to their location of origin and branch tip shape indicates whether it is a domesticated (square) or wild (circle) strain. New long-read genome assemblies presented in this study are indicated in bold. Major genomic elements found on mitochondria are shown and colored according to guide elements at the top. Inverted elements appear on the underside of the line. Duplicated elements are indicated with asterisk. Inset depicts untransformed phylogenetic tree with species labeled.

Previous analyses have suggested that hybridization events can generate discordance between species and mitochondrial phylogenies in yeast (De Chiara, et al. 2020; Peris, et al. 2017). To investigate this, we also included other *Saccharomyces* species with available long read data in our mitochondrial phylogenetic analyses. For the most part, our mitochondrial phylogenies matched our species-level phylogeny with the notable exception of a strain of *S. jurei* (NCYC3947), a recently described European species (Naseeb, et al. 2018) that appears to share mitochondrial ancestry with a subgroup of European strains of *S. paradoxus*. The mitochondrial genomes from this subgroup also contain large structural variations (previously described in Yue et al. 2017) not seen in other strains of *S. paradoxus* and *S. cerevisiae*, further supporting their shared ancestry (Fig. 5).

### Intraspecific spore viability

Lastly, we performed intraspecific crosses of each wild East Asian strain with the common lab strain Y55, to assess the level of reproductive isolation. As expected, we found a lower level of viable spores when crossing with a divergent wild strain as compared to self-crossing Y55 (ANOVA F(7,152) = 9.63, p < 0.001, **Fig. S24**). Most crosses with East Asian strains reduced spore viability by ^~^50%, while crosses with HN1 reduced viability by ^~^75%.

## Discussion

Comparative genomic analyses have provided clues about the origin of Brewer’s yeast and have suggested an out-of-China origin (Peter, et al. 2018). Here, we provide seven new, high quality long/short-read genomes of highly divergent wild *S. cerevisiae* strains recently isolated in Far East Asia. Phylogenomic analyses of the long-read assemblies agree with previous findings that the wild East Asian strains (CHN, Taiwanese) are basal relative to other *S. cerevisiae* strains (Duan, et al. 2018; Peter, et al. 2018) and, in the case of the CHN IX and Taiwanese clades, show considerable divergence (Fig. 1). In addition, we show that the CHN IX clade (represented here by JXXY16.1 and XXYS1.4) and the one strain representing the Taiwanese clade (EM14S01-3B), likely compose a single monophyletic group distinct from not only the other East Asian strains in our study but also all other strains of *S. cerevisiae* sequenced to date.

Our comprehensive analysis of structural variations (SVs) further elucidates the evolutionary history and intraspecific diversity of *S. cerevisiae*. Structural variations were identified for each strain pair revealing patterns of genomic divergence. There is a noteworthy pattern of higher amounts of SVs in wild East Asian strains, especially in the three strains within the Clade IX Complex. As a species, *S. cerevisiae* has been shown to accumulate balanced variations at a slower rate compared to *S. paradoxus* (Yue, et al. 2017). This is likely due to the different selection histories of these species. Many *S. cerevisiae* strains have long been associated with human activities where domestication, cross-breeding and admixture have resulted in largely mosaic genomes (Hyma and Fay 2013; Liti 2015; Liti, et al. 2009), whereas *S. paradoxus* strains are recently isolated, wild strains. Interestingly, we found that wild East Asian strains accumulated both structural variations at a high rate, more similar to rates normally seen in *S. paradoxus* (Fig. 3). It has been suggested that the geographical isolation of some *S. paradoxus* subpopulations may have favored quick fixation of structural rearrangements (Leducq, et al. 2016). We may be witnessing similar patterns in the wild East Asian *S. cerevisiae* strains.

In context of the seven previously assembled *S. cerevisiae* long-read genomes and those from *Saccharomyces* sensu stricto species, our results reveal other important aspects of yeast evolutionary genomic history. Not only do the phylogenetic patterns we describe reveal discrete boundaries between certain clade-levels in terms of TEs, indicating that transfer of persisting TEs between deep-rooted clades either through horizontal gene transfer or hybridization is rare (Fig. 4). They also give us context for the evolutionary history of these elements in their own right. Interestingly, we found that *Ty5*, a relatively rare retrotransposon with no previously known functional versions in *S. cerevisiae*, has retained functionality in the divergent East Asian Clade IX complex. Additionally, we found that *Ty2*, a TE suggested to be a recent introduction to *S. cerevisiae* via *S. mikatae* (Carr, et al. 2012; Liti, et al. 2005), is also present in the East Asian Clade IX complex. This indicates that this event occurred early in *S. cerevisiae* history, that the donor-donee relationship is reversed, that it happened multiply, or that this element was lost in *S. paradoxus* and other closely related species. With respect to the latter hypothesis, our genomic survey indicates numerous of losses of functional different *Ty* elements in various strains suggesting that *Ty* extinction within clades is probably not uncommon and that near complete loss of all traces of extinct elements can occur relatively rapidly (see for example *Ty4* and *Ty5* in Fig. 4).

In conclusion, we suggest that the divergence of the East Asian Clade IX Complex occurred prior to the genetically close-knit, global radiation of *S. cerevisiae* strains we see today, potentially before their domestication. This begs the question whether there are truly wild *S. cerevisiae* strains outside of Asia at all, especially if the colonization of the rest of the world happened contemporarily with humans. Overall, this study generates new, valuable genomic resources and expands our understanding of the genetic variation and evolutionary history of one of the most important organisms in human history, *S. cerevisiae*. Moreover this set of high-quality genomes, encompassing both domesticated and wild populations from different ecological backgrounds, provides an important resource for future explorations into the dynamics that govern eukaryotic genome evolution.

## Methods

### Yeast strain origins

We selected eight *Saccharomyces cerevisiae* strains for long-read sequencing and genome assembly (Table 1). Seven of these strains originate from East Asia. Six strains were isolated in China (Duan, et al. 2018; Wang, et al. 2012) from a variety of ecological niches and one in Taiwan (Peter, et al. 2018). The six Chinese strains cover many of the lineages (CHN I, II, IV, VI, and IX) previously shown to be highly divergent from other *S. cerevisiae* strains based on short-read sequencing. The final strain (Y55) is a common laboratory strain isolated in France with a known mosaic genomic background originating in West Africa. To place our analyses in context, we also included currently publicly available *Saccharomyces* sensu stricto long read genome assemblies as well as assemblies from *Torulaspora delbrueckii* and *Kluyveromyces lactis* (**Table S3**).

**Table 1:** Descriptions of the *Saccharomyces cerevisiae* strains sequenced in this study.

| Lineage | Strain | Species | Source | Geographic Location |
| --- | --- | --- | --- | --- |
| CHN I | HN1 | *S. cerevisiae* | rotten wood, primeval forest | Diaoluo Mountain, Hainan, China |
| CHN II | SX2 | *S. cerevisiae* | bark of a Fagaceae tree, primeval forest | Qinling Mountain, Shaanxi, China |
| CHN IV | HLJ1 | *S. cerevisiae* | bark of Quercus mongolica, secondary forest | Jingbo lake, Heilongjiang, China |
| CHN VI | BJ4 | *S. cerevisiae* | intestine of a butterfly, park | Haidian, Beijing, China |
| CHN IX | JXXY16.1 | *S. cerevisiae* | bark, primeval forest | Xiangxiyuan, Hubei Province, China |
| CHN IX | XXYS1.4 | *S. cerevisiae* | bark, primeval forest | Xiangxiyuan, Hubei Province, China |
| Taiwanese | EM14S01-3B | *S. cerevisiae* | soil | Taiwan |
| West African+ | Y55 | *S. cerevisiae* | wine grapes | France |

### DNA preparation and long-read sequencing

Before sequencing, strains were sporulated and tetrads were dissected to allow for autodiploidization, making strains homozygous across all loci. Strains were incubated at 30°C in 5ml YEPD (1% yeast extract, 2% peptone, 2% dextrose) in a shaking incubator for 24h before we harvested cells by centrifugation. We extracted genomic DNA using NucleoSpin Microbial DNA extraction kit according to the manufacturer’s instructions (Macherey-Nagel). Genomic DNA for strain Y55 was extracted independently using the QIAGEN Blood & Culture DNA Midi Kit. Samples were sequenced on PacBio Sequel and Sequel II platforms at the NGI/Uppsala Genome Center (Science for Life Laboratory, Sweden) and the University of Minnesota Sequencing Center (USA). In addition to these PacBio data, we also used publicly available paired-end Illumina sequence data previously generated for each strain (**Table S1**).

### Genome assembly and annotation

Nuclear contigs were assembled with Flye v2.8.1 (**Fig. S1**, default settings, est. genome size = 12.4Mbp) (Kolmogorov, et al. 2019). We used short read sequences for each strain to error-correct the long reads using FMLRC v2 (Wang, et al. 2018). Corrected long reads and the short reads were subsequently used to polish the Flye assemblies using Racon v1.4.13 (Vaser, et al. 2017) and POLCA v3.4.2 (Zimin and Salzberg 2019) respectively. We further scaffolded the contigs based on the reference S288C genome (GCA_000146045.2) using RaGOO v1.1 (Alonge, et al. 2019) and filled any gaps this generated using multiple iterations of LR Gapcloser v1 (Xu, et al. 2019) and Gapcloser (Luo, et al. 2012). To account for any errors introduced by using long reads to fill gaps, we further polished each assembly once more using Racon v1.4.13 and POLCA v3.4.2. Mitochondrial assemblies were largely assembled using Flye without the assumption of even coverage (--metagenomic) using all long reads as input. JXXY16.1 and Y55 mitochondrial genomes were assembled using Flye v2.8.1 with default settings. Mitochondrial contigs were extracted by mapping the Flye output to the reference mitochondrial genome using Nucmer (Delcher, et al. 2002). These assemblies were polished and scaffolded following the same process as that of the nuclear assemblies. Completeness of the final genome assemblies was assessed using BUSCO v4.0.5 (Simão, et al. 2015; Waterhouse, et al. 2018).

We annotated nuclear genes, mitochondrial genes, centromeres, transposable elements, core X elements and Y-prime elements using modified versions of the pipelines within the LRSDAY package (Yue and Liti 2018). In addition to our eight newly assembled genomes, we also used the same method to annotate the previously published long read assemblies (**Table S3**). Nuclear genes orthologous to annotated genes in the *S. cerevisiae* S288C reference genome were identified using Proteinortho v6.0.24 (Lechner, et al. 2011). Genes for which no orthologous protein was found in the reference were clustered based on orthology to each other.

To further characterize *Ty* elements, we determined potential element viability by translating coding regions of full elements based on reading frames identified for each element in S288C. Elements containing premature stop codons or extensive frameshifts were categorized as putatively being reproductively inviable (loss-of-function or LOF). Additionally, we created gene trees for whole elements of each *Ty* class using MAFFT v7.471 alignments (default settings) with PhyML v3.0 (substitution model = HKY85; bootstrap = 100; tree searching using SRT and NNI; conducted in Unipro UGENE v36.0). To determine the likelihood of closely related elements within a given strain resulting from transposition or segmental genome duplication, we mapped the 10000bp regions containing each element to related intra-strain elements.

### Phylogenomic analysis

To place our eight assembled genomes within the context of other *Saccharomyces* strains, we employed both a consensus gene tree and Assembly and Alignment-Free (AAF) approaches to phylogenetic tree construction. For consensus species trees, we used OrthoFinder v2.4.0 (Fig. 1) in addition to a standard gene tree approach. For the latter, we aligned all orthologous genes found in at least 5 strains (5847 genes) using MUSCLE v3.8.31 (Edgar 2004) and performed maximum-likelihood single-tree inference for each locus using RAxML-NG v1.01 (Kozlov et al. 2019) with a discrete GAMMA model of rate heterogeneity. We used Astral-III v5.7.4 (Zhang et al. 2018) with these gene trees to generate a consensus species tree.

AAF v20171001 (Fan, et al. 2015) was used with a k-mer size of 20 nucleotides and a threshold frequency of 7 for each k-mer to be included in the analysis. AAF was used to compare the long-read and short-read sequencing data for the 25 *Saccharomyces* strains (**Fig. S2, Table S3**). Short-read sequencing data for *K. lactis* and *T. delbrueckii* were included as outgroups.

To generate the mitochondrial phylogeny, we reoriented the start of each assembly based on the position of the tRNA gene, trnP(ugg), then aligned these assemblies to each other using Mugsy v.1.2.3 (Angiuoli and Salzberg 2011). This multiple sequence alignment was then used to create a maximum likelihood tree using IQ-TREE v2.0.5 (options: -m TPM2u+F+R3 -B 1000 -bnni -alrt 1000) (Hoang, et al. 2018; Nguyen, et al. 2015). The model was determined using the ModelFinder component of IQ-Tree (Kalyaanamoorthy, et al. 2017).

### Structural variation detection

To identify the structural variations (SVs) between strains within *S. cerevisiae*, we performed exhaustive pairwise comparisons between the 15 strains with long-read assemblies (210 comparisons). We focused on five types of SV: deletions, insertions, tandem duplications, inversions and translocations. The SVs were detected using MUM&Co (O’Donnell and Fischer 2020), which utilizes MUMmer v3.23 (Marçais, et al. 2018) to perform whole-genome alignments and detect SVs >= 50bps.

### Spore viability assay

To assess the level of reproductive isolation between the divergent East Asian strains and modern *S. cerevisiae*, we crossed all strains with Y55 (α; ho; leu2Δ::HygMX) (and Y55 to itself) and assessed the spore viability of each cross. We sporulated each strain by incubating them in liquid sporulation medium (KAC; 2% potassium acetate) for 3 days at 23C. These cultures were then incubated with 10*μ*l zymolyase (100 U/ml) at 37C for 30min before being plated on YEPD (2.5% agar) in equal mixture with cultures of Y55 (α; ho; leu2Δ::HygMX) and grown for 48h at 30C. This culture was streaked on YEPD + hygromycin and replica plated to minimal media. A single colony was selected from each cross and grown up in liquid YEPD overnight, spun down, put in KAC, and incubated at room temperature with shaking for four days to induce sporulation. The resulting tetrads were treated with zymolyase for 30min at room temperature. 500*μ*l of sterile water were added before spores were dissected out of the tetrads onto YPD plates, using a Singer MSM 400 micro-manipulator. We dissected 20 tetrads yielding 80 spores per cross. Plates were incubated at 30C and colonies were counted after 72h, indicating viable spores that were able to germinate.

Respiratory competence was determined by plating strains of yeast on rich media containing nonfermentable glycerol as the sole carbon source (1% yeast extract, 2% peptone, 2% glycerol).

## Data Availability Statement

The data underlying this article are available in the European Nucleotide Archive and can be accessed with accession number **PRJEB38713**.

## Acknowledgments

This work was supported by grants from Stockholm University (Science for Life Laboratory sequencing grant SU FV-2.2.2-1843-17 to RS), the Swedish Research Council (2017-04963 to RS), the Knut and Alice Wallenberg Foundation (2017.0163 to RS), the Carl Trygger foundation (CTS 17: 431 to ZZ), the Wenner-Gren Foundations (UPD2018-0196, UPD2019-0110 to DPB), and The University of Minnesota Department of Ecology, Evolution, and Behavior. We acknowledge the support of the National Genomics Infrastructure (NGI)/Uppsala Genome Center and UPPMAX for providing assistance in massive parallel sequencing and computational infrastructure. Work performed at NGI/Uppsala Genome Center (project SNIC 2019/8-23) has been funded by RFI/VR and Science for Life Laboratory, Sweden. We would like to thank Feng-Yan Bai and Gianni Liti for donating strains, Jia-Xing Yue for advice on the LRSDAY pipeline, and Gianni Liti, Samuel O’Donnell and Chris Wheat for discussion.

