## Supplementary Material for "Genomic evidence of an early evolutionary divergence event in wild *Saccharomyces cerevisiae*"

### Table S1: Metrics for nuclear and mitochondrial assemblies

| | Metric | Y55 | BJ4 | HLJ1 | HN1 | SX2 | EM14S01-3B | JXXY16.1 | XXYS1.4 |
|---|---|---|---|---|---|---|---|---|---|
| Nuclear | **Mean read depth** | 99.65 | 165.91 | 45.31 | 66.14 | 174.18 | 76.84 | 25.96 | 76.46 |
| | **Total sequence count** | 16 | 16 | 16 | 16 | 16 | 16 | 16 | 16 |
| | **Total sequence length (bp)** | 11791742 | 11888108 | 11702338 | 11820137 | 11918758 | 11803938 | 11692380 | 11654158 |
| | **Min sequence length (bp)** | 230382 | 225758 | 229169 | 236734 | 226383 | 209147 | 223390 | 199212 |
| | **Max sequence length (bp)** | 1479087 | 1489054 | 1458639 | 1481187 | 1479650 | 1470812 | 1471675 | 1483244 |
| | **Mean sequence length (bp)** | 736983.88 | 743006.75 | 731396.12 | 738758.56 | 744922.38 | 737746.12 | 730773.75 | 728384.88 |
| | **Median sequence length (bp)** | 741368.5 | 752361.5 | 741606.5 | 758822 | 748579.5 | 755211 | 741203.5 | 733903.5 |
| | **N50 (bp)** | 916013 | 918555 | 895159 | 913524 | 928563 | 897577 | 902407 | 894632 |
| | **L50** | 6 | 6 | 6 | 6 | 6 | 6 | 6 | 6 |
| | **N90 (bp)** | 454735 | 426239 | 433662 | 434566 | 433099 | 444590 | 425488 | 432873 |
| | **L90** | 13 | 13 | 13 | 13 | 13 | 13 | 13 | 13 |
| | **GC%** | 38% | 38% | 38% | 38% | 38% | 38% | 38% | 38% |
| | **N%** | 0% | 0% | 0% | 0% | 0% | 0% | 0% | 0% |
| | **Genome size (Mbp)** | 11.79 | 11.89 | 11.7 | 11.82 | 11.92 | 11.8 | 11.69 | 11.65 |
| Mitochondria | **Total sequence count** | 1 | 1 | 1 | 1 | 1 | 1 | 1 | 1 |
| | **Total sequence length (bp)** | 80847 | 86123 | 78472 | 88103 | 80393 | 86728 | 68736 | 85245 |
| | **GC%** | 17% | 15% | 15% | 14% | 15% | 15% | 15% | 15% |
| | **N%** | 0% | 0% | 0% | 0% | 0% | 0% | 0% | 0% |

### Table S2: Results of BUSCO genome completion analysis

| Metric | Y55 | BJ4 | HLJ1 | HN1 | SX2 | EM14S01-3B | JXXY16.1 | XXYS1.4 |
|---|---|---|---|---|---|---|---|---|
| **Complete BUSCOs** | 98.7 | 98.5 | 98.2 | 98.4 | 98.5 | 98.5 | 97.6 | 96.5 |
| **Complete and single-copy BUSCOs** | 96.8 | 96.7 | 96.4 | 96.7 | 96.7 | 96.7 | 95.8 | 94.9 |
| **Complete and duplicated BUSCOs** | 1.9 | 1.8 | 1.8 | 1.7 | 1.8 | 1.8 | 1.8 | 1.6 |
| **Fragmented BUSCOs** | 0.1 | 0.1 | 0.2 | 0.1 | 0.2 | 0.1 | 0.3 | 0.9 |
| **Missing BUSCOs** | 1.2 | 1.4 | 1.6 | 1.5 | 1.3 | 1.4 | 2.1 | 2.6 |
| **Total BUSCO groups searched** | 2137 | 2137 | 2137 | 2137 | 2137 | 2137 | 2137 | 2137 |

### Table S3: Information about long and short-read sequencing datasets used in this study.

| Genome | Species | Long-read Sequencing | | | Short-read Sequencing | | |
|---|---|---|---|---|---|---|---|
| | | Source | Instrument | ENA Accession | Source | Instrument | ENA Accession |
| HN1 | S. cerevisiae | **this study** | PacBio Sequel | PRJEB38713 | Duan et al. 2018 | Illumina HiSeq 2000 | PRJNA396809 |
| SX2 | S. cerevisiae | **this study** | PacBio Sequel | PRJEB38713 | Duan et al. 2018 | Illumina HiSeq 2000 | PRJNA396809 |
| HLJ1 | S. cerevisiae | **this study** | PacBio Sequel | PRJEB38713 | Duan et al. 2018 | Illumina HiSeq 2000 | PRJNA396809 |
| BJ4 | S. cerevisiae | **this study** | PacBio Sequel | PRJEB38713 | Duan et al. 2018 | Illumina HiSeq 2000 | PRJNA396809 |
| JXXY16.1 | S. cerevisiae | **this study** | PacBio Sequel | PRJEB38713 | Duan et al. 2018 | Illumina HiSeq 2000 | PRJNA396809 |
| XXYS1.4 | S. cerevisiae | **this study** | PacBio Sequel | PRJEB38713 | Duan et al. 2018 | Illumina HiSeq 2000 | PRJNA396809 |
| EM14S01-3B | S. cerevisiae | **this study** | PacBio Sequel | PRJEB38713 | Peter et al. 2018 | Illumina HiSeq 2000 | PRJEB13017 |
| Y55 | S. cerevisiae | **this study** | PacBioSequel | PRJEB38713 | NA | Illumina HiSeq 4000 | PRJNA552112 |
| S288C | S. cerevisiae | Yue et al. 2017 | PacBio RS | PRJEB7245 | Yue et al. 2017 | Illumina HiSeq 2500 | PRJNA340312 |
| DBVPG6044 | S. cerevisiae | Yue et al. 2017 | PacBio RS | PRJEB7245 | Yue et al. 2017 | Illumina HiSeq 2500 | PRJNA340312 |
| DBVPG6765 | S. cerevisiae | Yue et al. 2017 | PacBio RS | PRJEB7245 | Yue et al. 2017 | Illumina HiSeq 2500 | PRJNA340312 |
| SK1 | S. cerevisiae | Yue et al. 2017 | PacBio RS | PRJEB7245 | Yue et al. 2017 | Illumina HiSeq 2500 | PRJNA340312 |
| Y12 | S. cerevisiae | Yue et al. 2017 | PacBio RS | PRJEB7245 | Yue et al. 2017 | Illumina HiSeq 2500 | PRJNA340312 |
| YPS128 | S. cerevisiae | Yue et al. 2017 | PacBio RS | PRJEB7245 | Yue et al. 2017 | Illumina HiSeq 2500 | PRJNA340312 |
| UWOPS03-461.4 | S. cerevisiae | Yue et al. 2017 | PacBio RS | PRJEB7245 | Yue et al. 2017 | Illumina HiSeq 2500 | PRJNA340312 |
| CBS432 | S. paradoxus | Yue et al. 2017 | PacBio RS | PRJEB7245 | Yue et al. 2017 | Illumina HiSeq 2500 | PRJNA340312 |
| N44 | S. paradoxus | Yue et al. 2017 | PacBio RS | PRJEB7245 | Yue et al. 2017 | Illumina HiSeq 2500 | PRJNA340312 |
| YPS138 | S. paradoxus | Yue et al. 2017 | PacBio RS | PRJEB7245 | Yue et al. 2017 | Illumina HiSeq 2500 | PRJNA340312 |
| UFRJ50816 | S. paradoxus | Yue et al. 2017 | PacBio RS | PRJEB7245 | Yue et al. 2017 | Illumina HiSeq 2500 | PRJNA340312 |
| UWOPS91-917.1 | S. paradoxus | Yue et al. 2017 | PacBio RS | PRJEB7245 | Yue et al. 2017 | Illumina HiSeq 2500 | PRJNA340312 |
| NCYC3947 | S. jurei | Naseeb et al. 2018 | PacBio RS II | PRJEB24816 | Naseeb et al. 2018 | Illumina HiSeq 2500 | PRJEB24816 |
| CR85 | S. kudriavzevii | Macías et al. 2019 | Roche 454 | PRJEB31099 | Boonekamp et al. 2018 | Illumina HiSeq 2500 | PRJNA480800 |
| CBS12357 | S. eubayanus | NA | Nanopore MinION | PRJNA450912 | Hebly et al. 2015 | Illumina HiSeq 2500 | PRJNA450912 |
| COFT1 | T. delbrueckii | NA | Nanopore MinION | PRJNA435462 | Gomez-Angulo et al. 2015 | Genome Analyzer IIx | PRJNA278774 |
| GG799 | K. lactis | NA | PacBio | PRJNA386037 | NA | NA | NA |

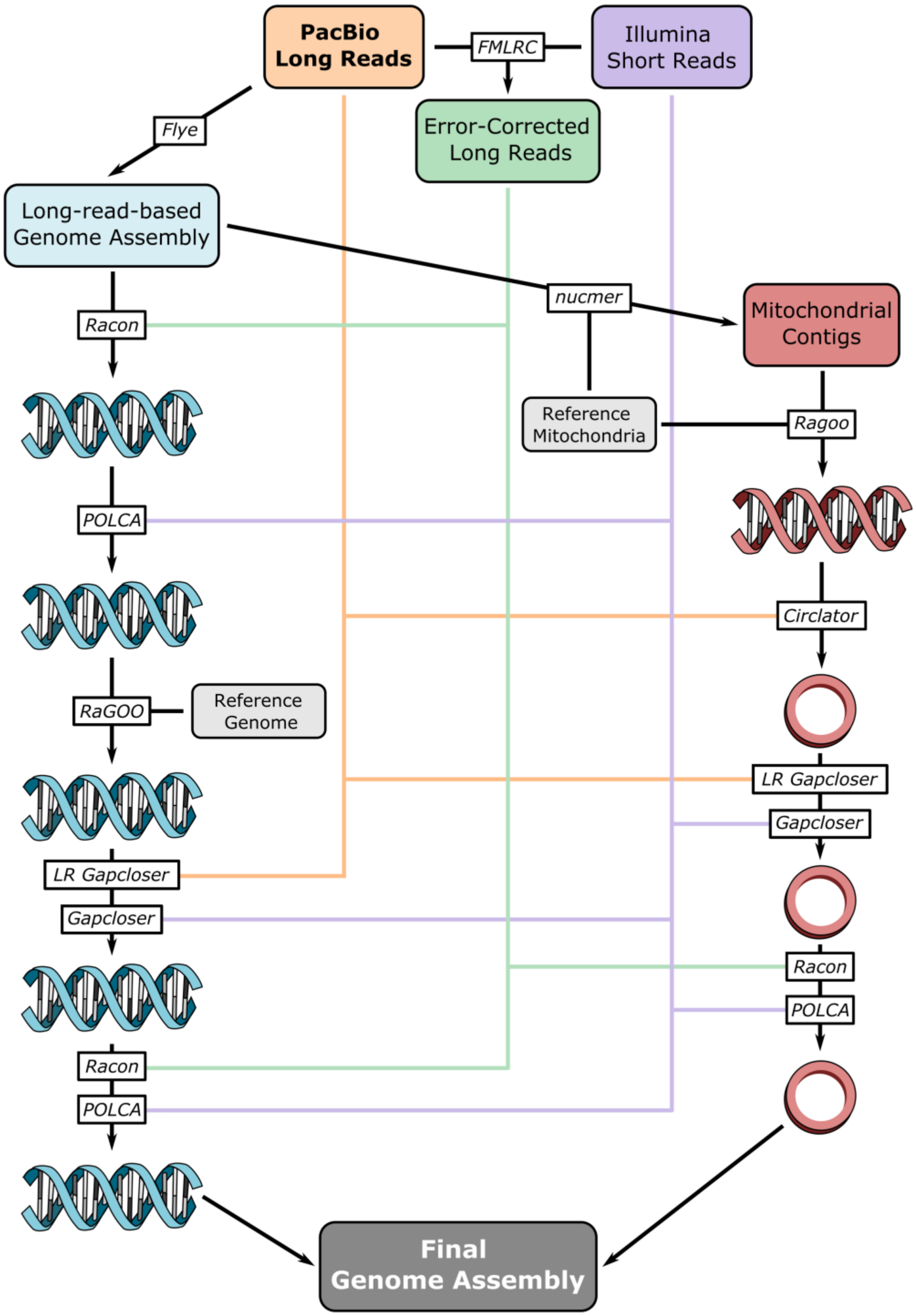


**Figure S1: Nuclear and mitochondrial genome assembly pipeline.** Workflow for assembly of nuclear (blue) and mitochondrial (red) genomes. PacBio long reads generated in this study are indicated in orange, while publicly available short-reads (**Table S3**) are indicated in purple. It is important to note that the Flye assembly processes were independent for nuclear and mitochondrial assembly, as indicated in the methods. The reference nuclear and mitochondrial genome used for assembly was S288C.

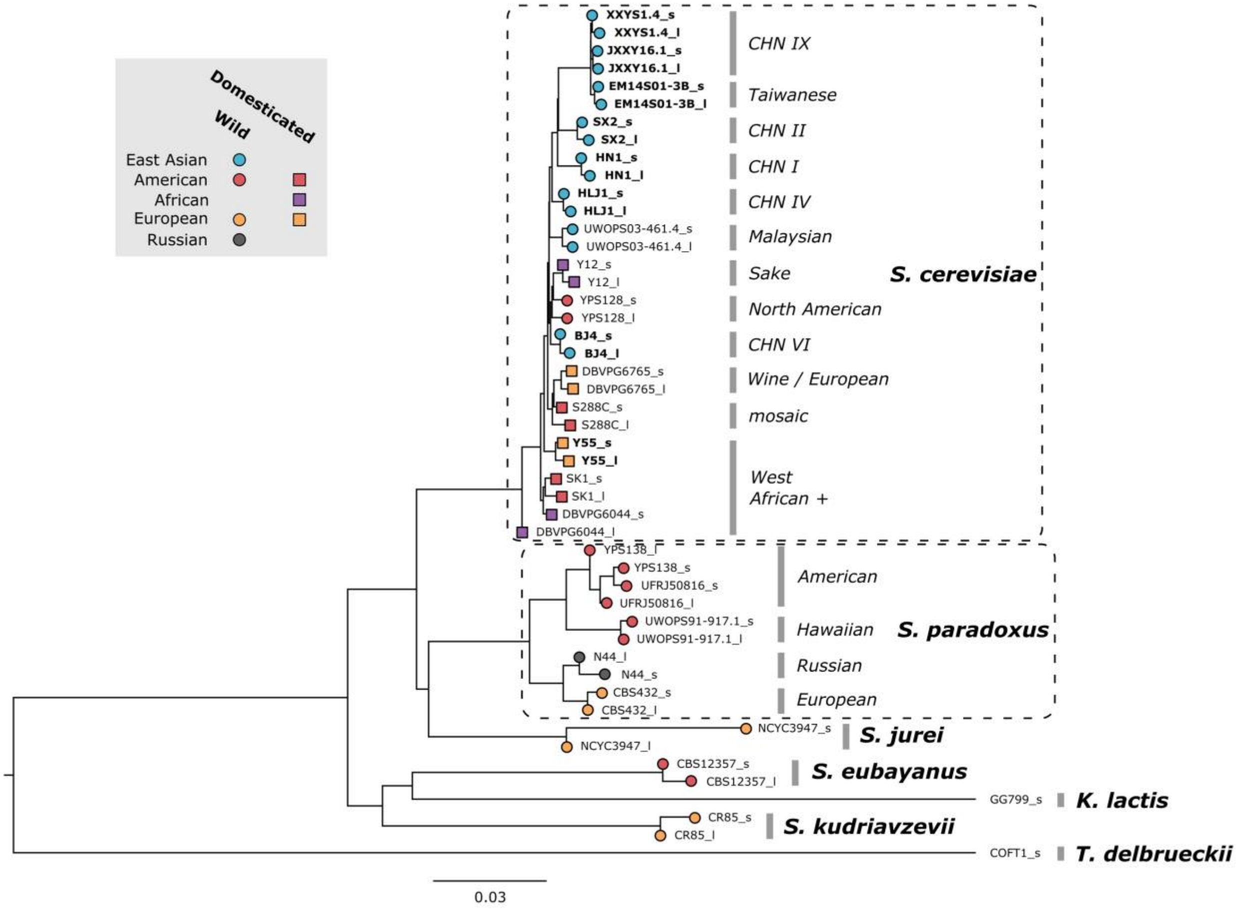


**Figure S2: Alignment and Assembly-Free (AAF) phylogenetic network of short and long-read sequencing data** Phylogenetic network generated using an Alignment and Assembly-Free (AAF) approach. The analysis was performed with k-mer size of 20 and a threshold frequency of 7 for each k-mer to be included in the analysis. For species with more than a single long-read genome assembly (*S. cerevisiae and S. paradoxus*), species clades are indicated in italics. *Saccharomyces* strains are colored according to their location of origin and branch tip shape indicates whether it is a domesticated (square) or wild (circle) strain. New long-read sequencing data presented in this study are indicated in bold. Short-read and long-read sequencing data are labeled '_s' and '_l' respectively, following the strain name.

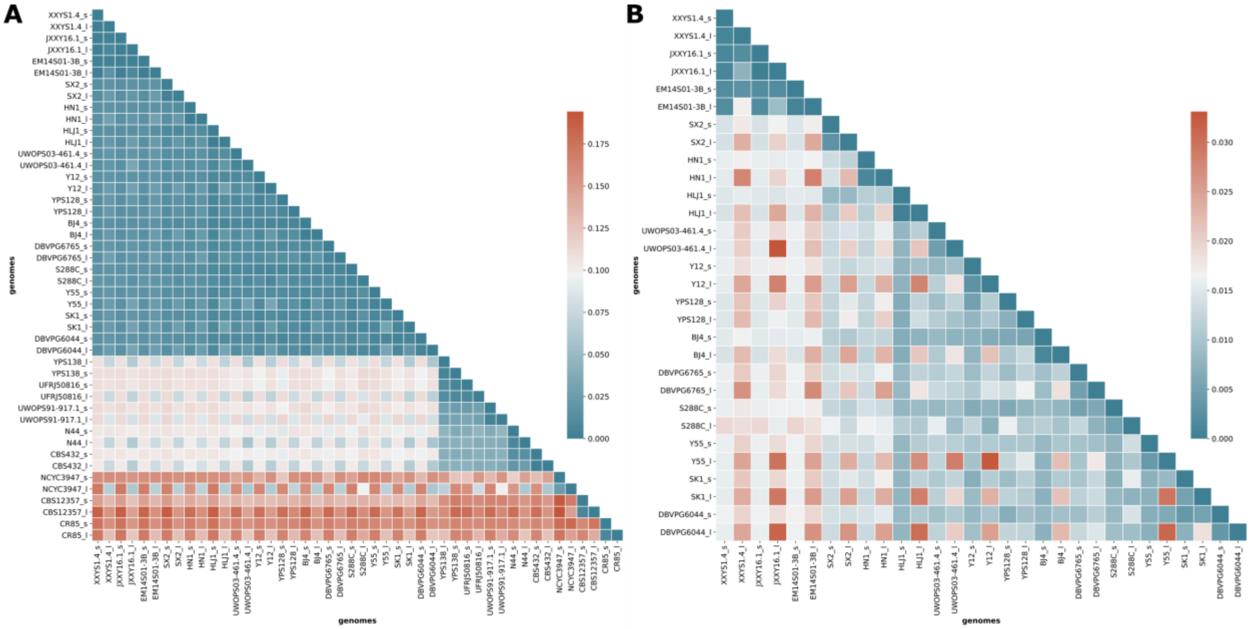


**Figure S3: Assembly and Alignment Free (AAF) Distance matrix of short-read vs long-read (A)** Pairwise distance matrix generated from AAF of short read and long-read sequencing data. Short-read and long-read sequencing data are labeled '_s' and '_l' respectively, following the strain name. The analysis was performed with k-mer size of 20 and a threshold frequency of 7 for each k-mer to be included in the analysis. **(B)** A closer look at only *S. cerevisiae* strains.

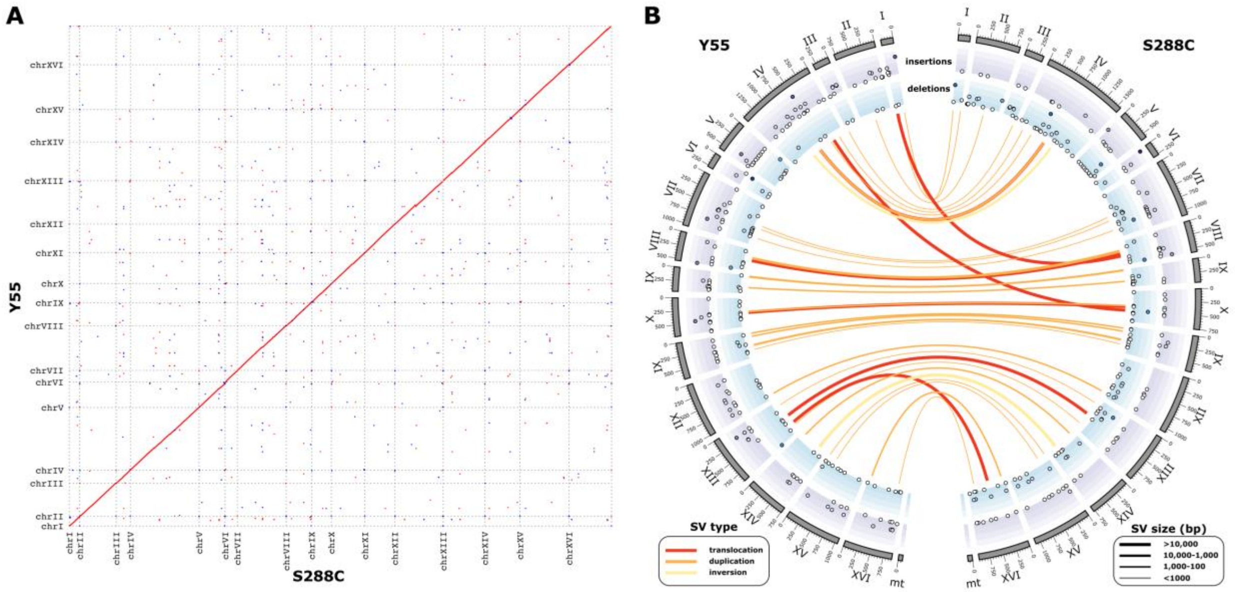


**Figure S4: Y55 genome assembly (A)** Genome comparison of the reference strain, S288C, and Y55. Sequence homology within the dot plots is indicated by red dots for forward matches and blue dots for reverse matches. **(B)** CIRCOS plot showing the detected structural variations between reference strain, S288C and Y55. Translocations (red), duplications (orange) and inversions (yellow) are depicted as links between the two genomes. The width of the link reflects the relative size of the variation (bp). Insertions (blue) and deletions (purple) are depicted in the outer tracks. Deletion and insertion size increase towards the outside. Chromosome size is depicted on the outside in 1kb units.

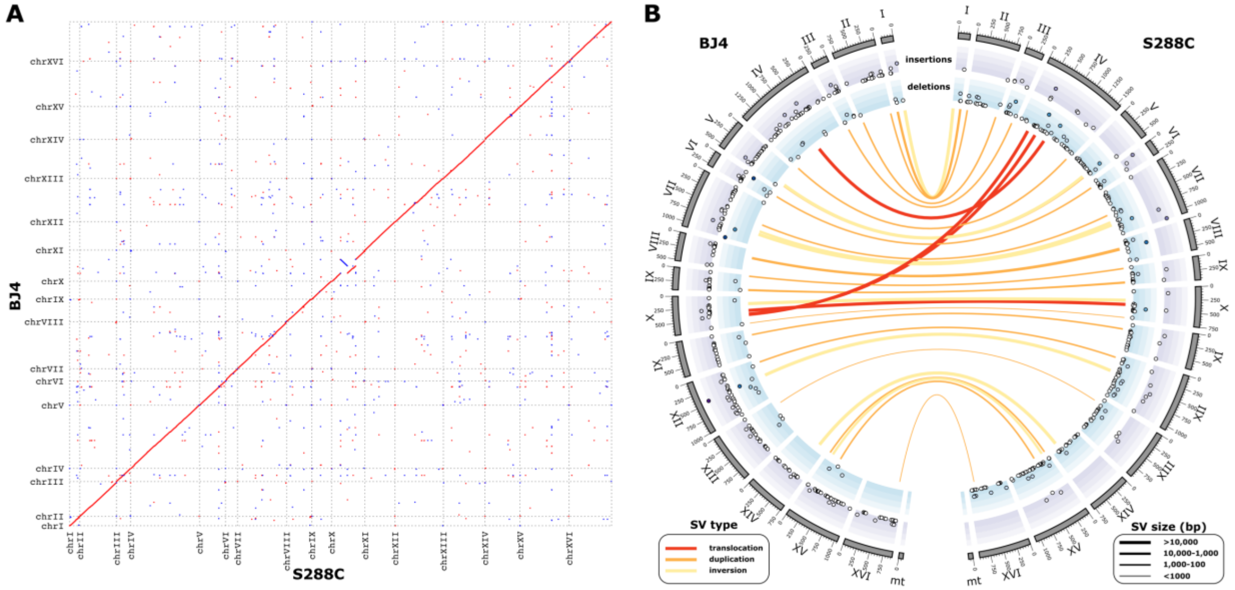


**Figure S5: BJ4 genome assembly (A)** Genome comparison of the reference strain, S288C, and BJ4. Sequence homology within the dot plots is indicated by red dots for forward matches and blue dots for reverse matches. **(B)** CIRCOS plot showing the detected structural variations between reference strain, S288C and BJ4. Translocations (red), duplications (orange) and inversions (yellow) are depicted as links between the two genomes. The width of the link reflects the relative size of the variation (bp). Insertions (blue) and deletions (purple) are depicted in the outer tracks. Deletion and insertion size increase towards the outside. Chromosome size is depicted on the outside in 1kb units.

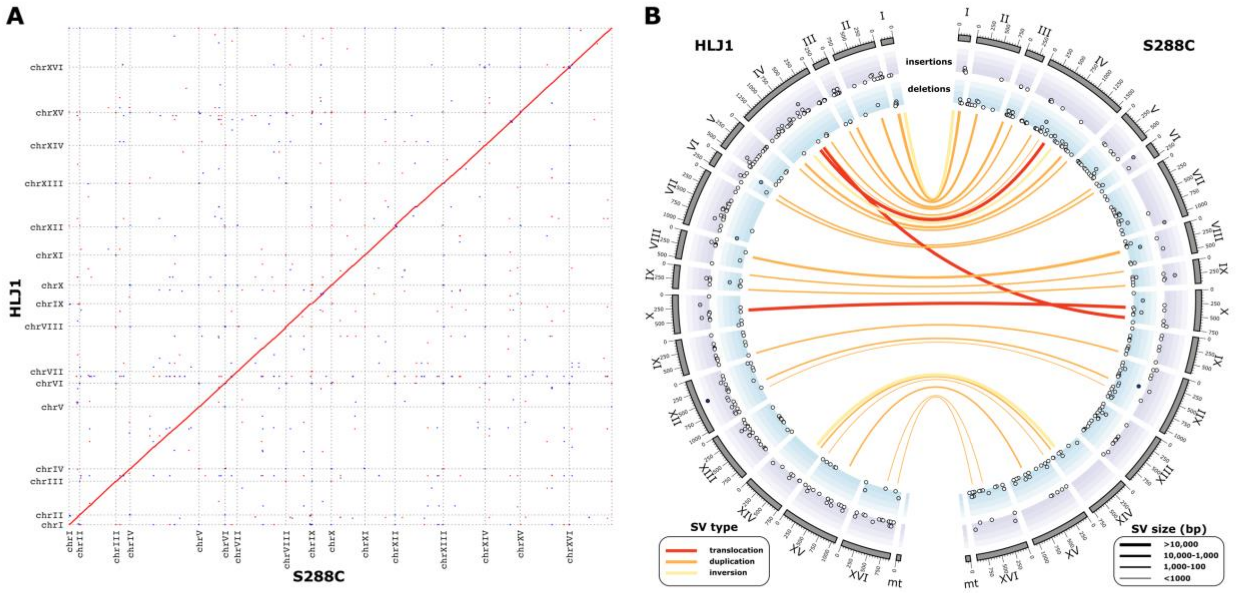


**Figure S6: HLJ1 genome assembly (A)** Genome comparison of the reference strain, S288C, and HLJ1. Sequence homology within the dot plots is indicated by red dots for forward matches and blue dots for reverse matches. **(B)** CIRCOS plot showing the detected structural variations between reference strain, S288C and HLJ1. Translocations (red), duplications (orange) and inversions (yellow) are depicted as links between the two genomes. The width of the link reflects the relative size of the variation (bp). Insertions (blue) and deletions (purple) are depicted in the outer tracks. Deletion and insertion size increase towards the outside. Chromosome size is depicted on the outside in 1kb units

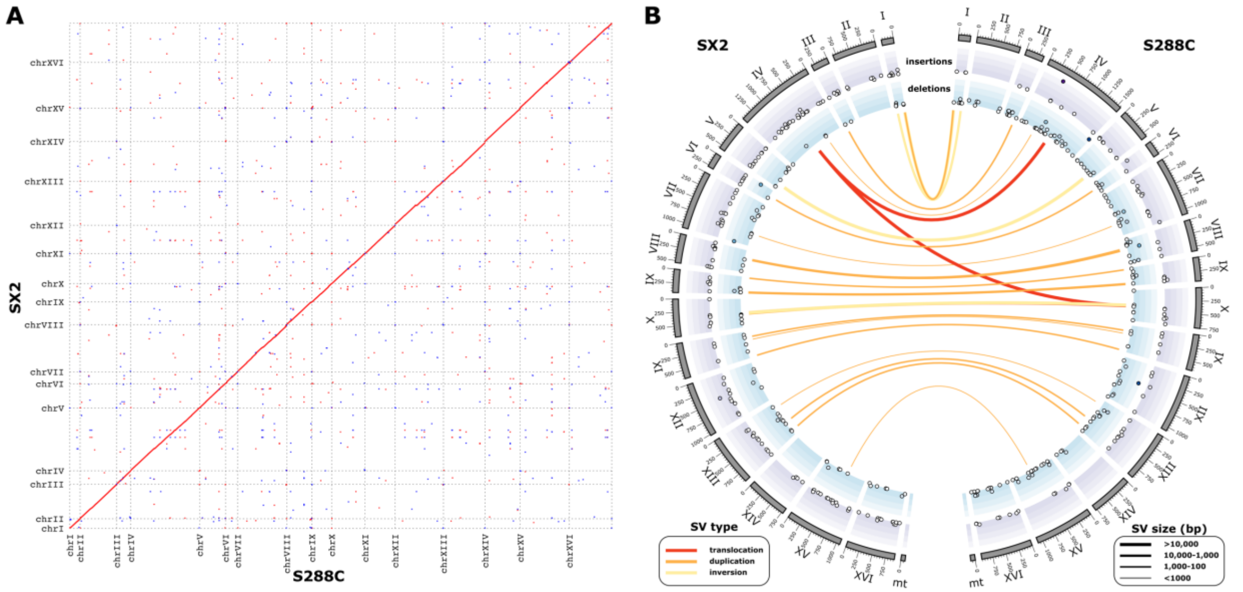


**Figure S7: SX2 genome assembly (A)** Genome comparison of the reference strain, S288C, and SX2. Sequence homology within the dot plots is indicated by red dots for forward matches and blue dots for reverse matches. **(B)** CIRCOS plot showing the detected structural variations between reference strain, S288C and SX2. Translocations (red), duplications (orange) and inversions (yellow) are depicted as links between the two genomes. The width of the link reflects the relative size of the variation (bp). Insertions (blue) and deletions (purple) are depicted in the outer tracks. Deletion and insertion size increase towards the outside. Chromosome size is depicted on the outside in 1kb units.

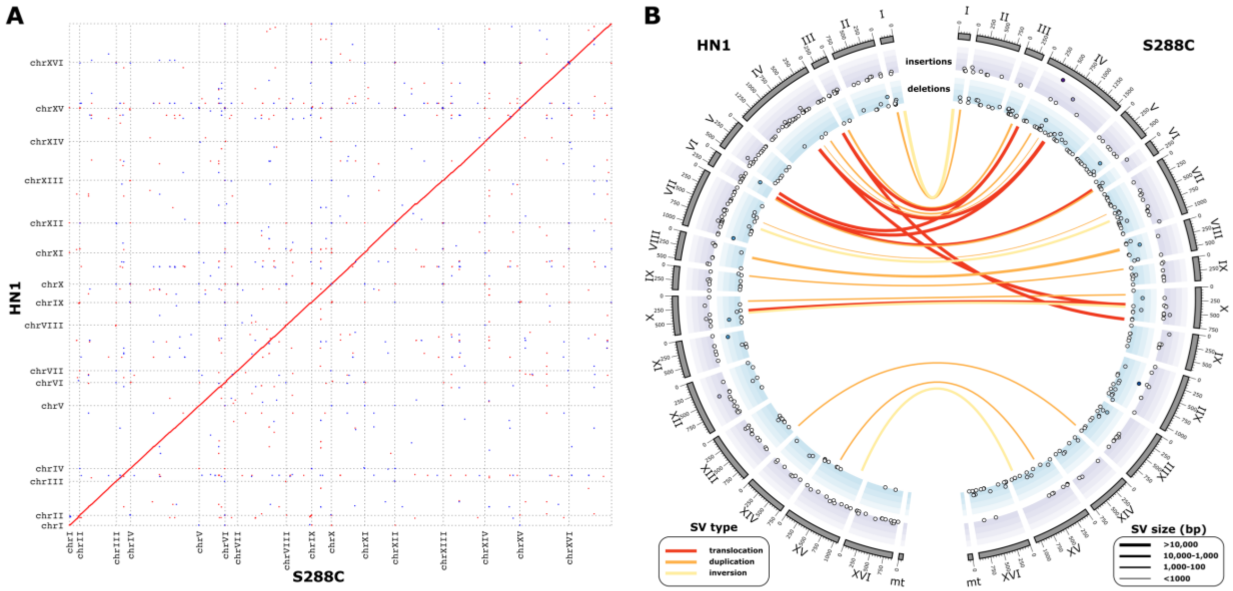


**Figure S8: HN1 genome assembly (A)** Genome comparison of the reference strain, S288C, and HN1. Sequence homology within the dot plots is indicated by red dots for forward matches and blue dots for reverse matches. **(B)** CIRCOS plot showing the detected structural variations between reference strain, S288C and HN1. Translocations (red), duplications (orange) and inversions (yellow) are depicted as links between the two genomes. The width of the link reflects the relative size of the variation (bp). Insertions (blue) and deletions (purple) are depicted in the outer tracks. Deletion and insertion size increase towards the outside. Chromosome size is depicted on the outside in 1kb units.

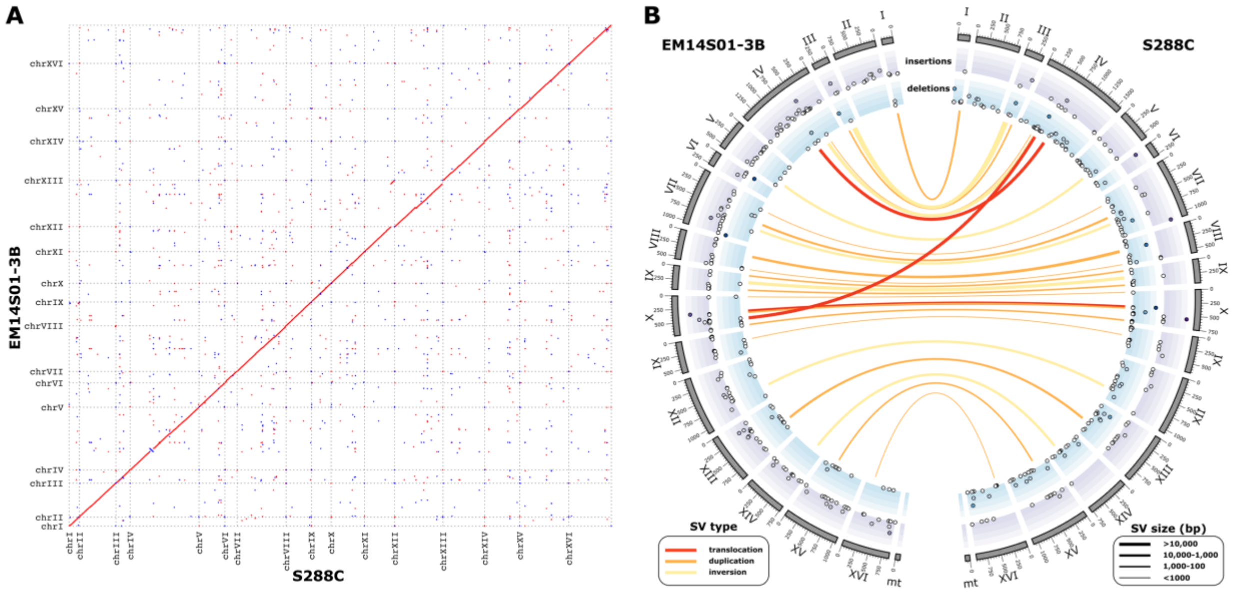


**Figure S9: EM14S01-3B genome assembly (A)** Genome comparison of the reference strain, S288C, and EM14S01-3B. Sequence homology within the dot plots is indicated by red dots for forward matches and blue dots for reverse matches. **(B)** CIRCOS plot showing the detected structural variations between reference strain, S288C and EM14S01-3B. Translocations (red), duplications (orange) and inversions (yellow) are depicted as links between the two genomes. The width of the link reflects the relative size of the variation (bp). Insertions (blue) and deletions (purple) are depicted in the outer tracks. Deletion and insertion size increase towards the outside. Chromosome size is depicted on the outside in 1kb units.

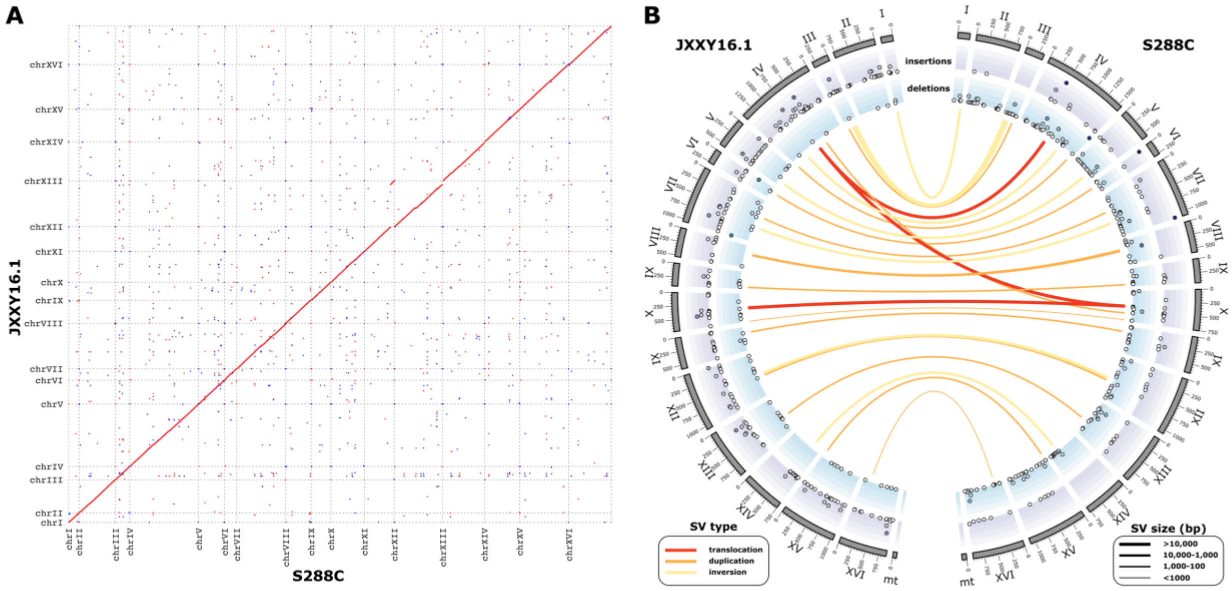


**Figure S10: JXXY16.1 genome assembly (A)** Genome comparison of the reference strain, S288C, and JXXY16.1. Sequence homology within the dot plots is indicated by red dots for forward matches and blue dots for reverse matches. **(B)** CIRCOS plot showing the detected structural variations between reference strain, S288C and JXXY16.1. Translocations (red), duplications (orange) and inversions (yellow) are depicted as links between the two genomes. The width of the link reflects the relative size of the variation (bp). Insertions (blue) and deletions (purple) are depicted in the outer tracks. Deletion and insertion size increase towards the outside. Chromosome size is depicted on the outside in 1kb units.

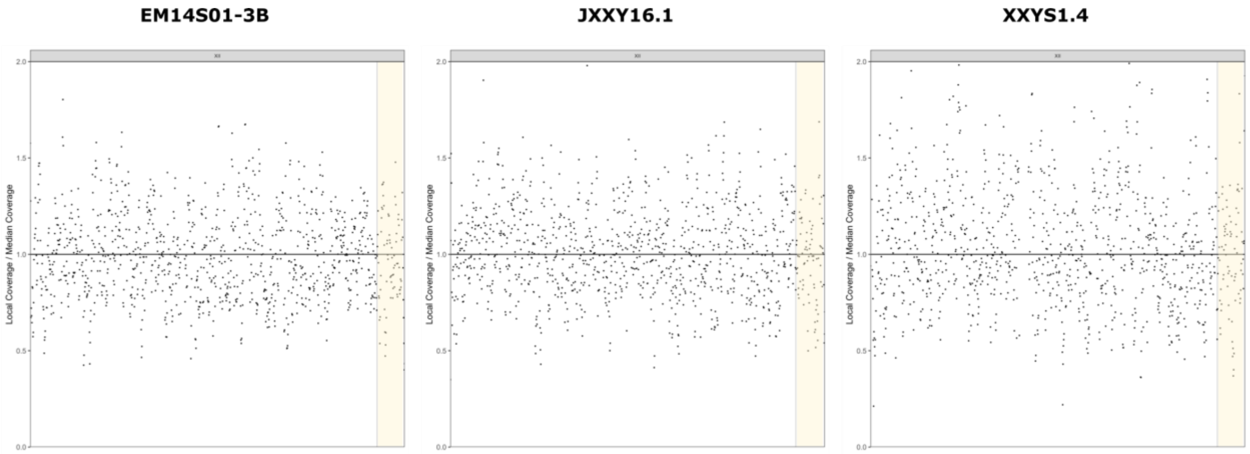


**Figure S11: Long-read sequencing coverage of chromosome XII in East Asian Clade IX strains.** X-axis indicates the length of the chromosome. Y-axis indicates the binned local coverage/ median coverage. Highlighted yellow region indicates the portion of the chromosome translocated from chromosome XI.

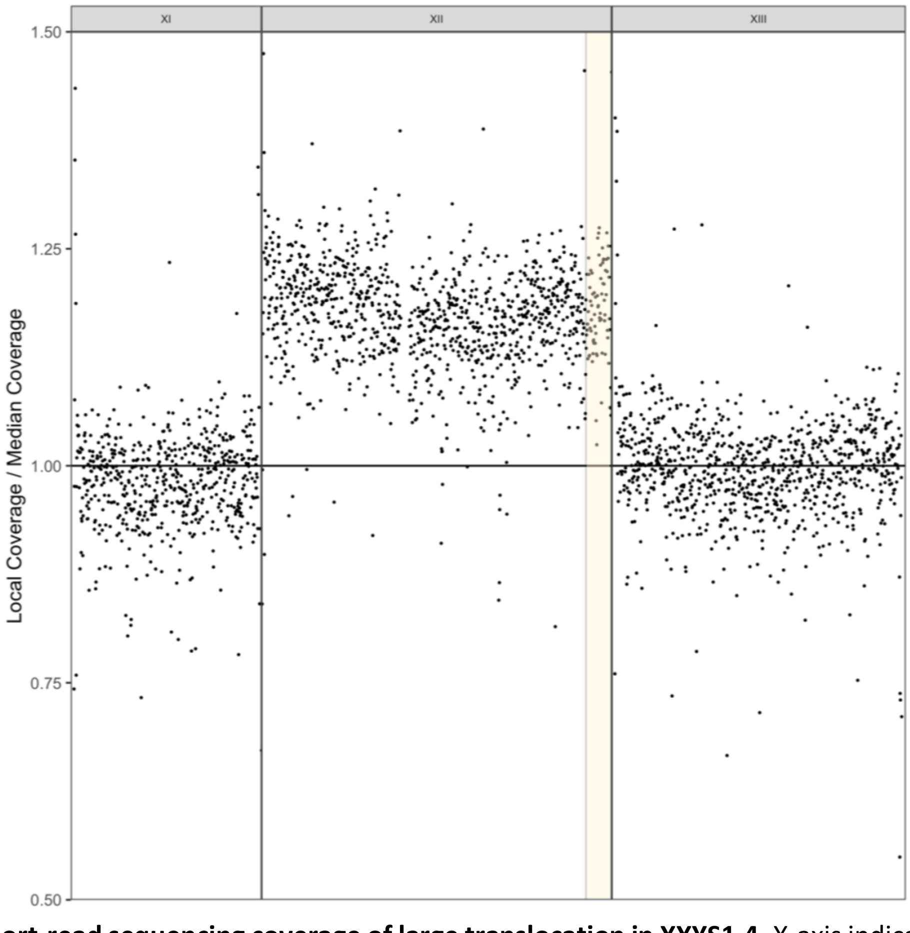


**Figure S12: Short-read sequencing coverage of large translocation in XXYS1.4.** X-axis indicates the length of the chromosome. Y-axis indicates the binned local coverage/ median coverage. Highlighted yellow region indicates the portion of the chromosome translocated from chromosome XI.

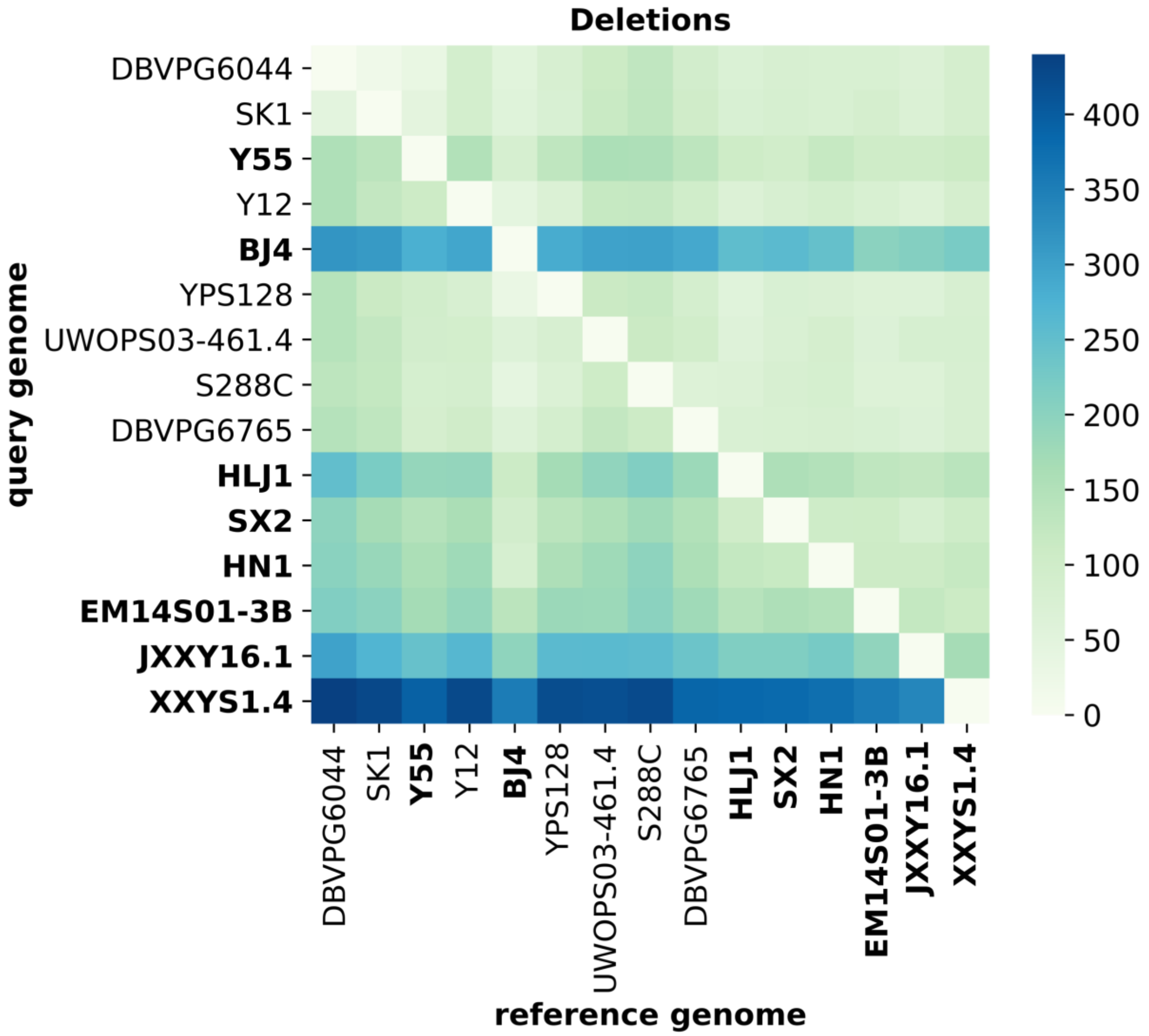


**Figure S13: Deletions within *Saccharomyces cerevisiae*.** Pairwise comparisons among all *Saccharomyces cerevisiae* genome assemblies depicting the total number of variations. Order of genome assemblies is consistent with the species tree (**Fig. 1**). New long-read genome assemblies presented in this study are indicated in bold.

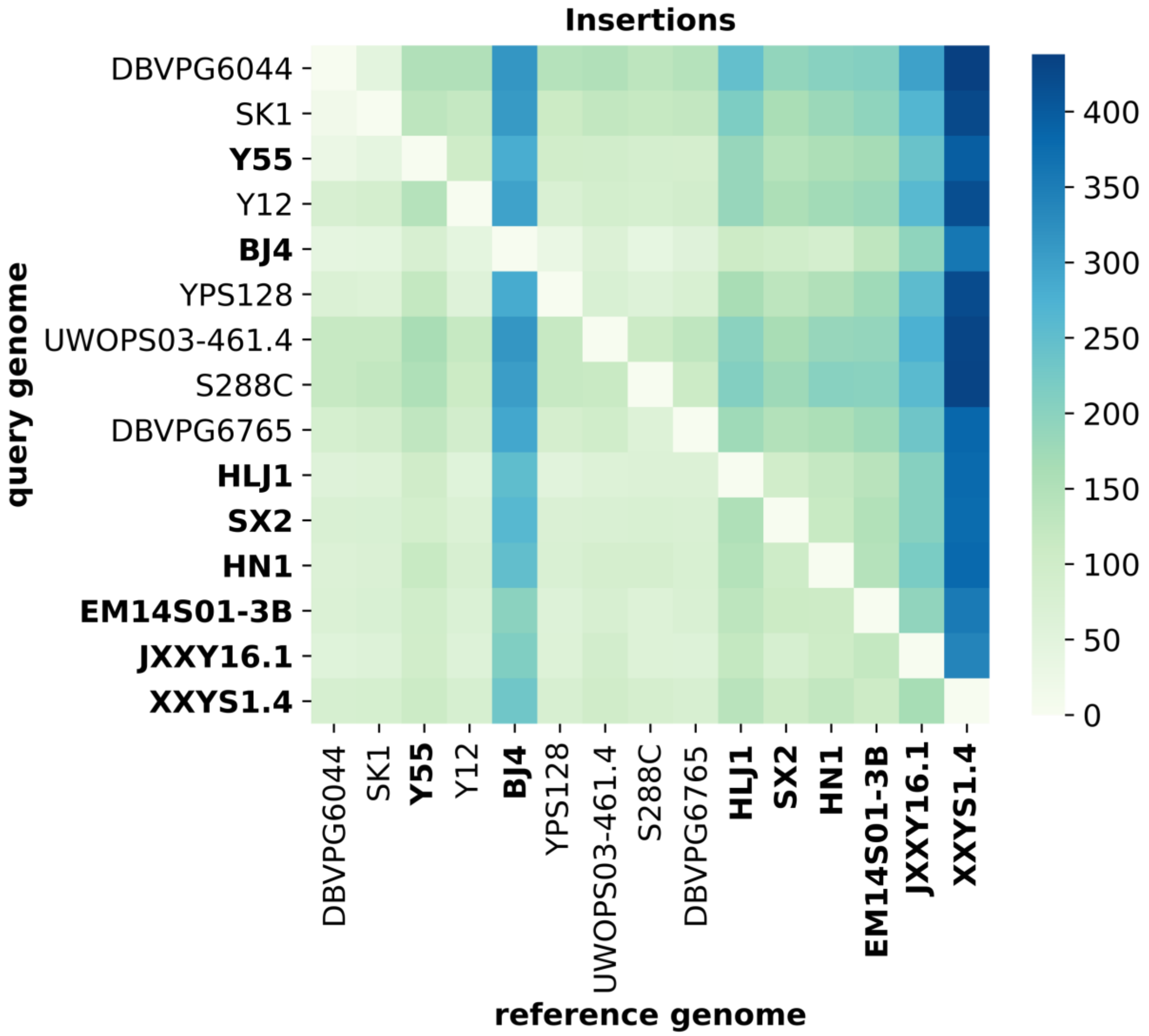


**Figure S14: Insertions within *Saccharomyces cerevisiae*.** Pairwise comparisons among all *Saccharomyces cerevisiae* genome assemblies depicting the total number of variations. Order of genome assemblies is consistent with the species tree (**Fig. 1**). New long-read genome assemblies presented in this study are indicated in bold.

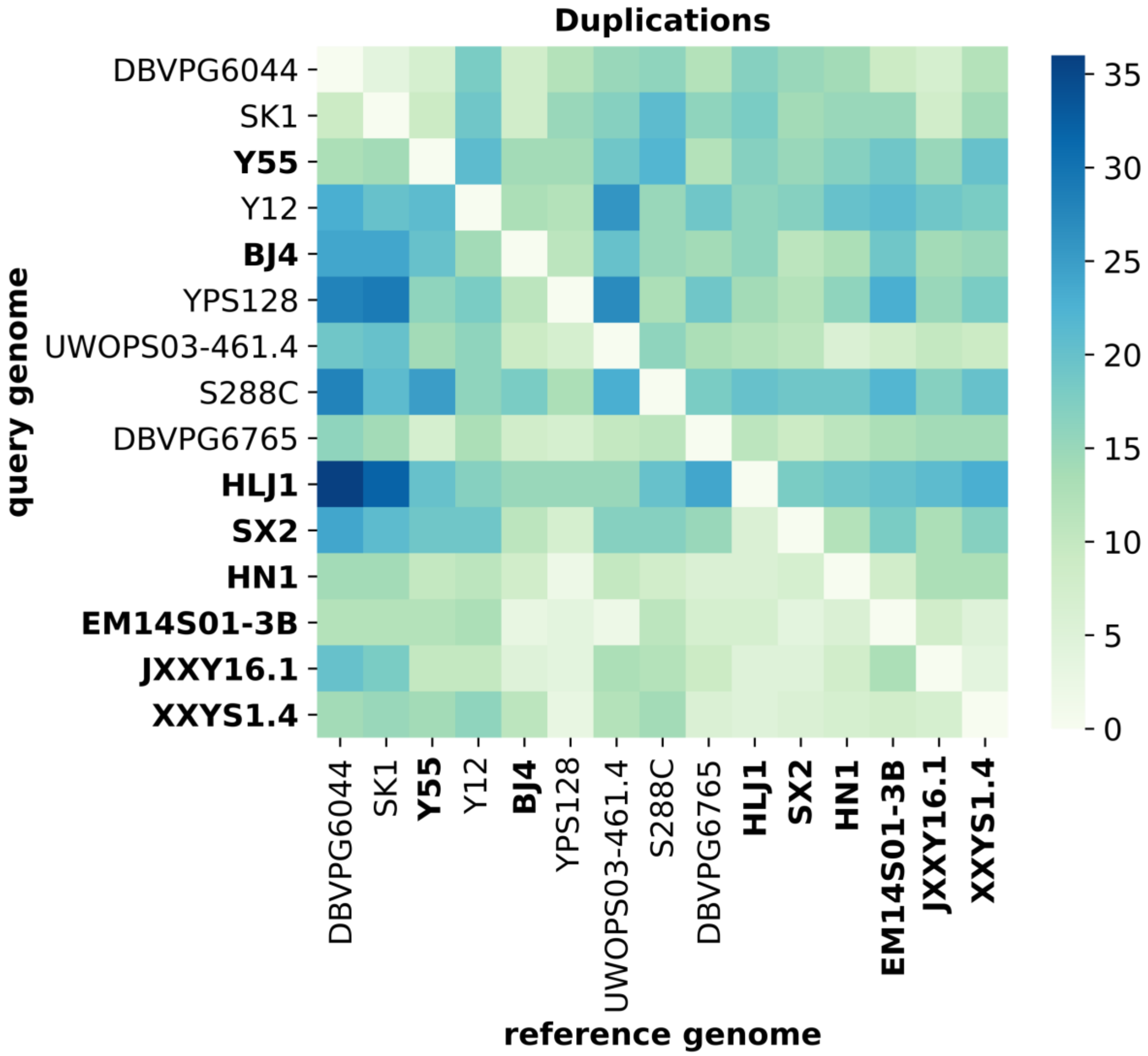


**Figure S15: Duplications within *Saccharomyces cerevisiae*.** Pairwise comparisons among all *Saccharomyces cerevisiae* genome assemblies depicting the total number of variations. Order of genome assemblies is consistent with the species tree (**Fig. 1**). New long-read genome assemblies presented in this study are indicated in bold.

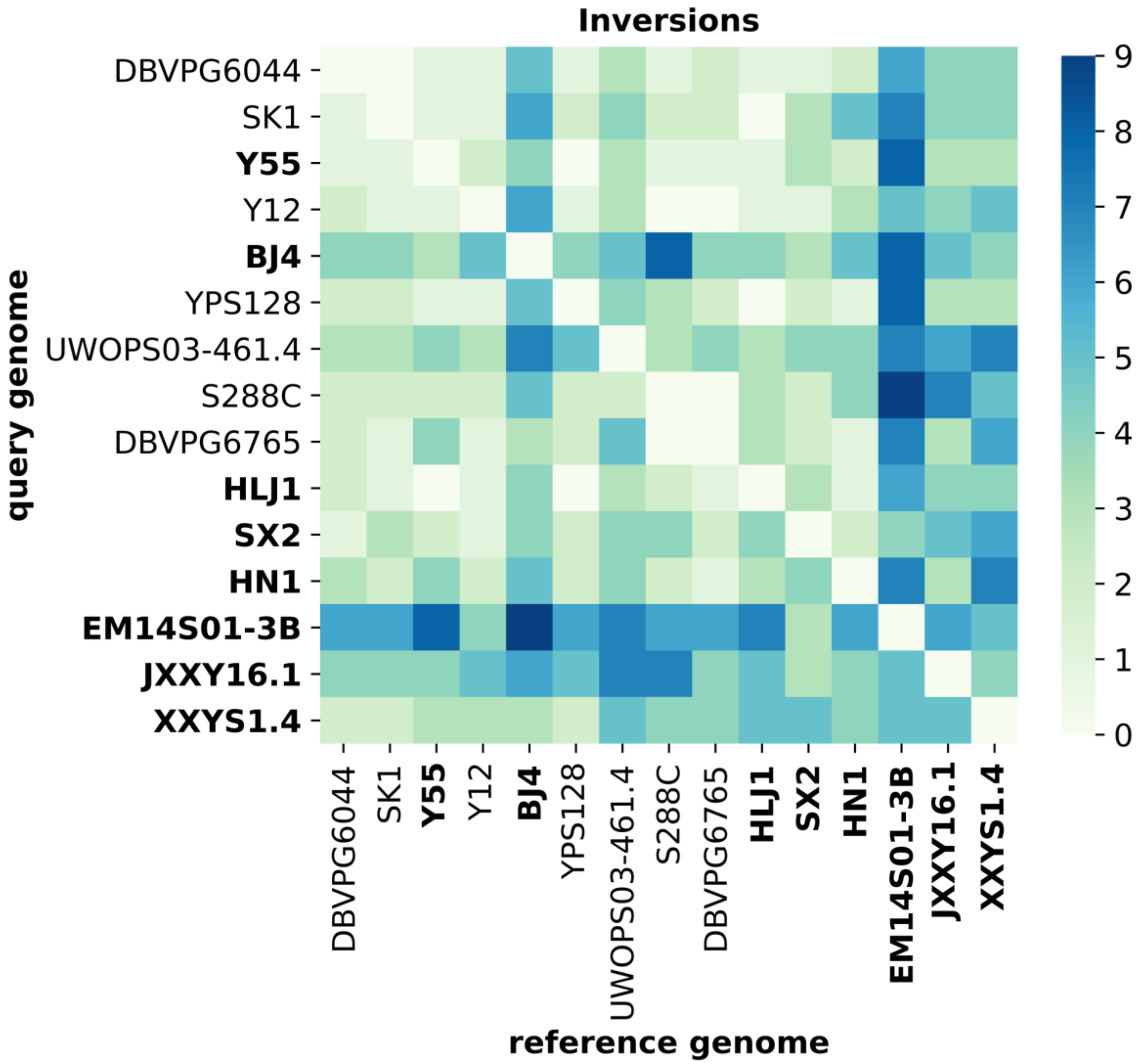


**Figure S16: Inversions within *Saccharomyces cerevisiae*.** Pairwise comparisons among all *Saccharomyces cerevisiae* genome assemblies depicting the total number of variations. Order of genome assemblies is consistent with the species tree (**Fig. 1**). New long-read genome assemblies presented in this study are indicated in bold.

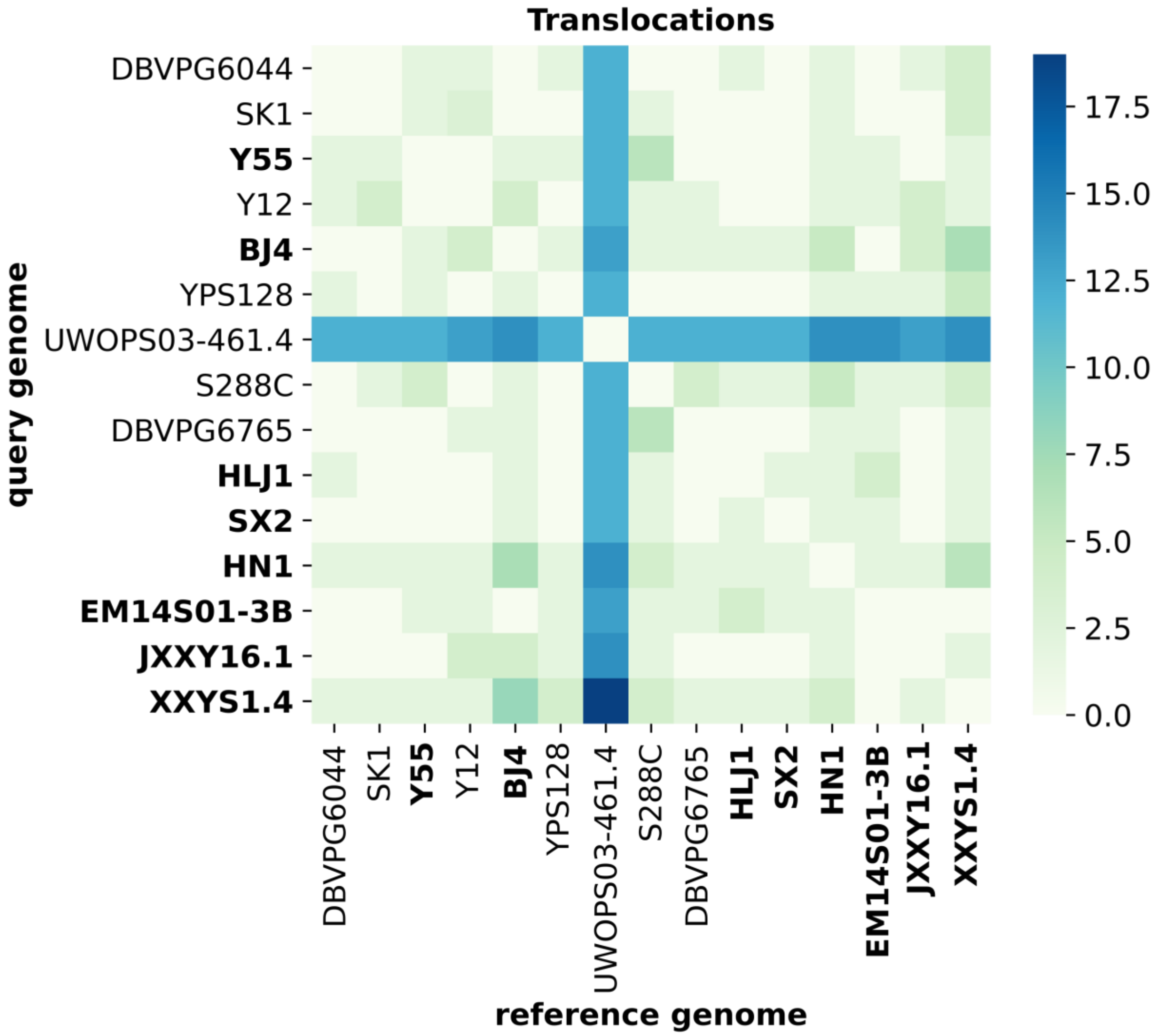


**Figure S17: Translocations within *Saccharomyces cerevisiae*.** Pairwise comparisons among all *Saccharomyces cerevisiae* genome assemblies depicting the total number of variations. Order of genome assemblies is consistent with the species tree (**Fig. 1**). New long-read genome assemblies presented in this study are indicated in bold.

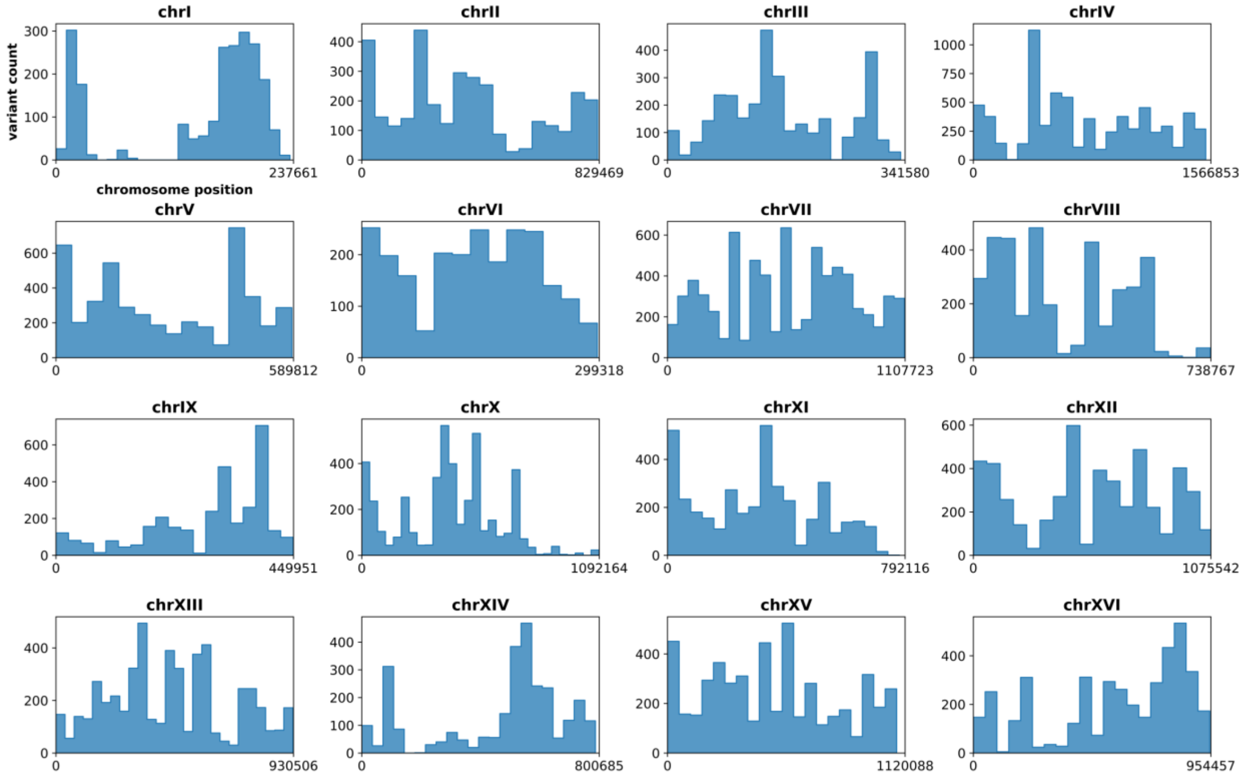


**Figure S18: Distribution of structural variations along chromosomes.** Histograms depicting the distribution of structural variations for each chromosome. Structural variations from all pairwise comparisons were included.

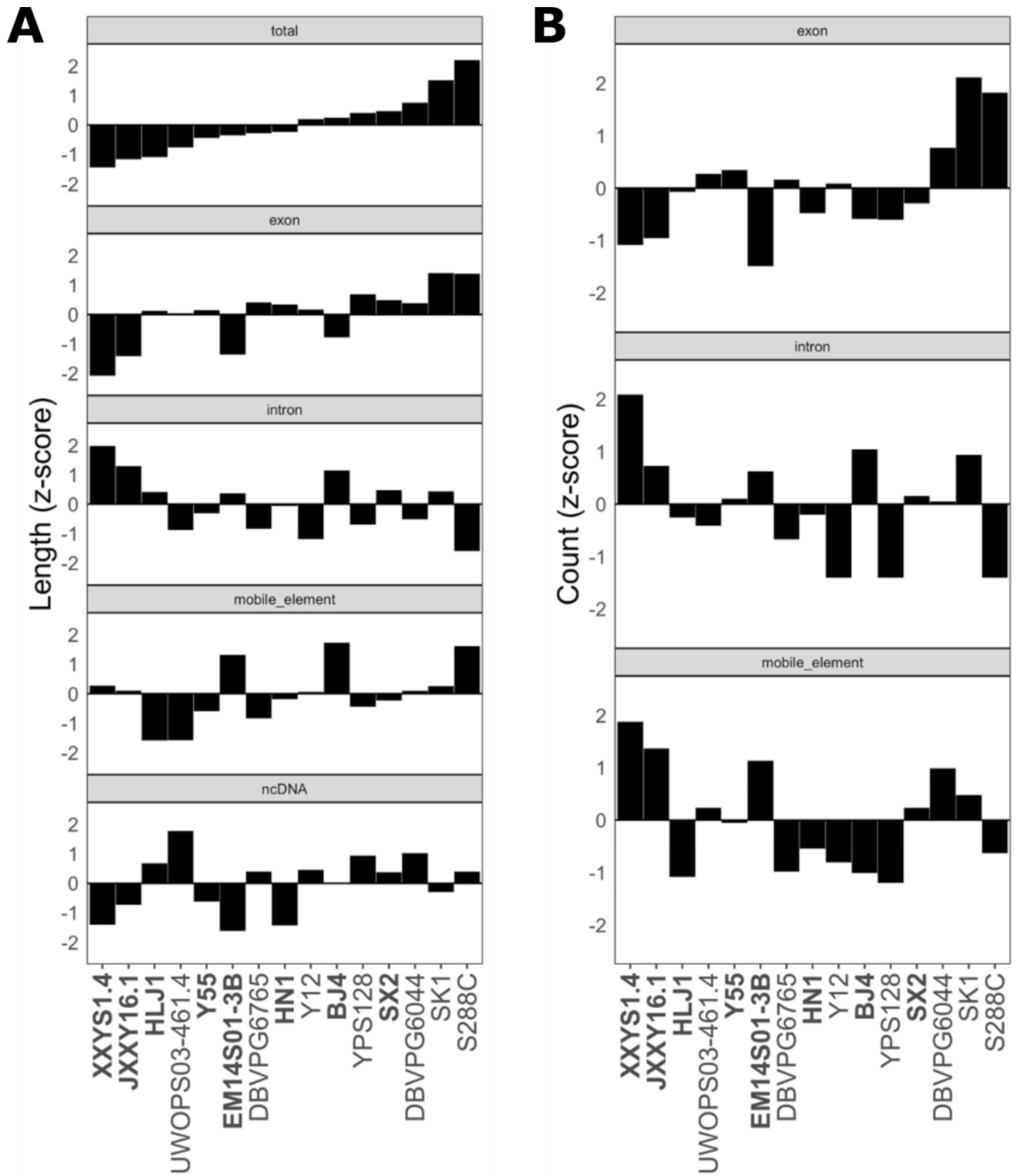


**Figure S19: Nuclear genome content (A)** Composite lengths in base pairs of genomic features relative to overall feature means for *S. cerevisiae* for total genomic length ($\bar{x}$=11.85Mbp, s=0.13Mbp), exon regions ($\bar{x}$=8.35 Mbp, s= 0.78 Mbp), intron regions ($\bar{x}$=79.27 kbp, s=5.27 kbp), and transposable elements (TEs) ($\bar{x}$=218.58 kbp, s=84.90 kbp). **(B)** Total counts of genomic features relative to overall feature means for *S. cerevisiae* for exon regions ($\bar{x}$=6104.07, s=54.31), intron regions ($\bar{x}$=79.93, s=19.16), and TEs ($\bar{x}$= 249.73, s= 68.85). Strains are ordered according to the total genomic length. Strains sequenced for this study are in bold.

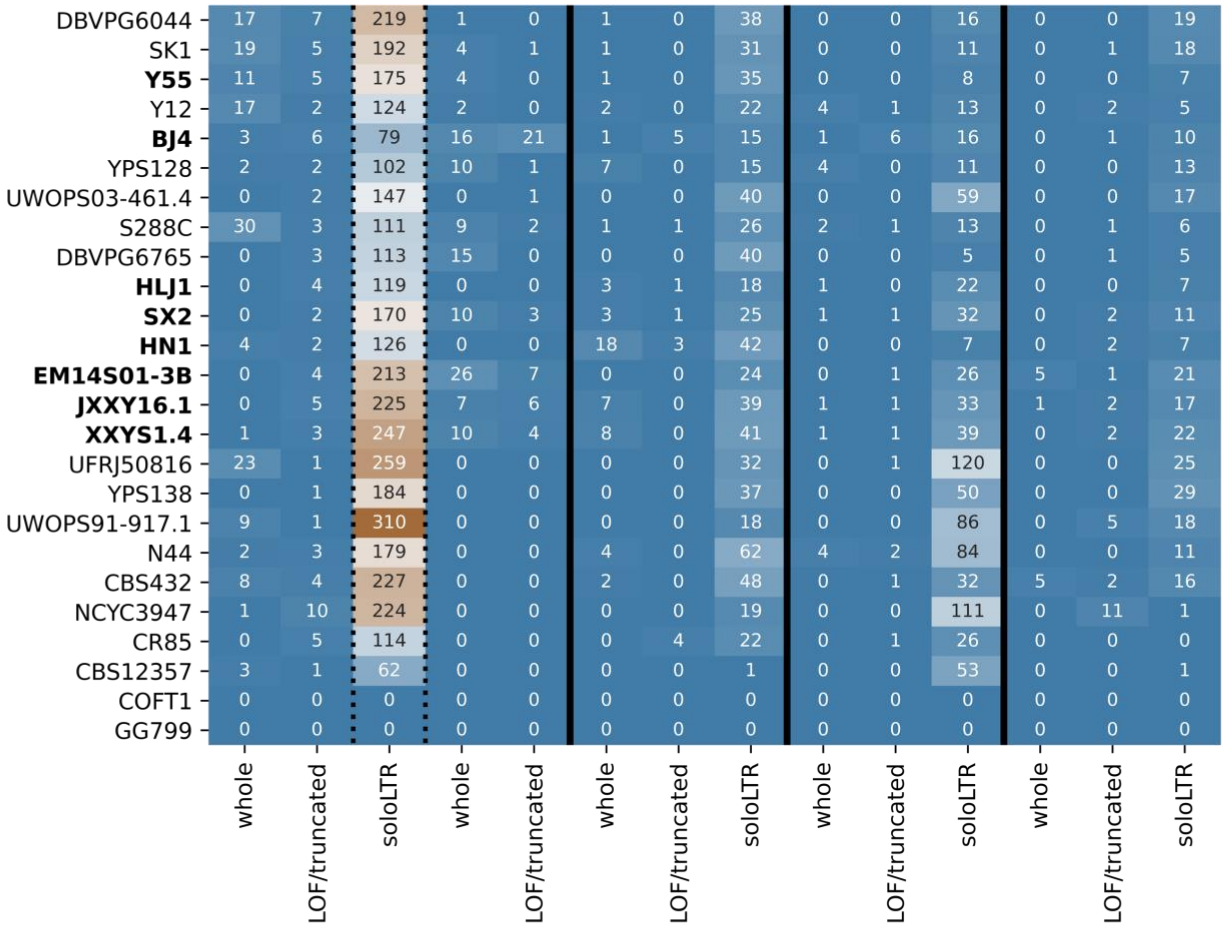


**Figure S20: Annotated transposable element composition in Saccharomyces** Total length of each type of transposable element found within each strain. Directly corresponds to the counts shown in **Fig. 3A**.

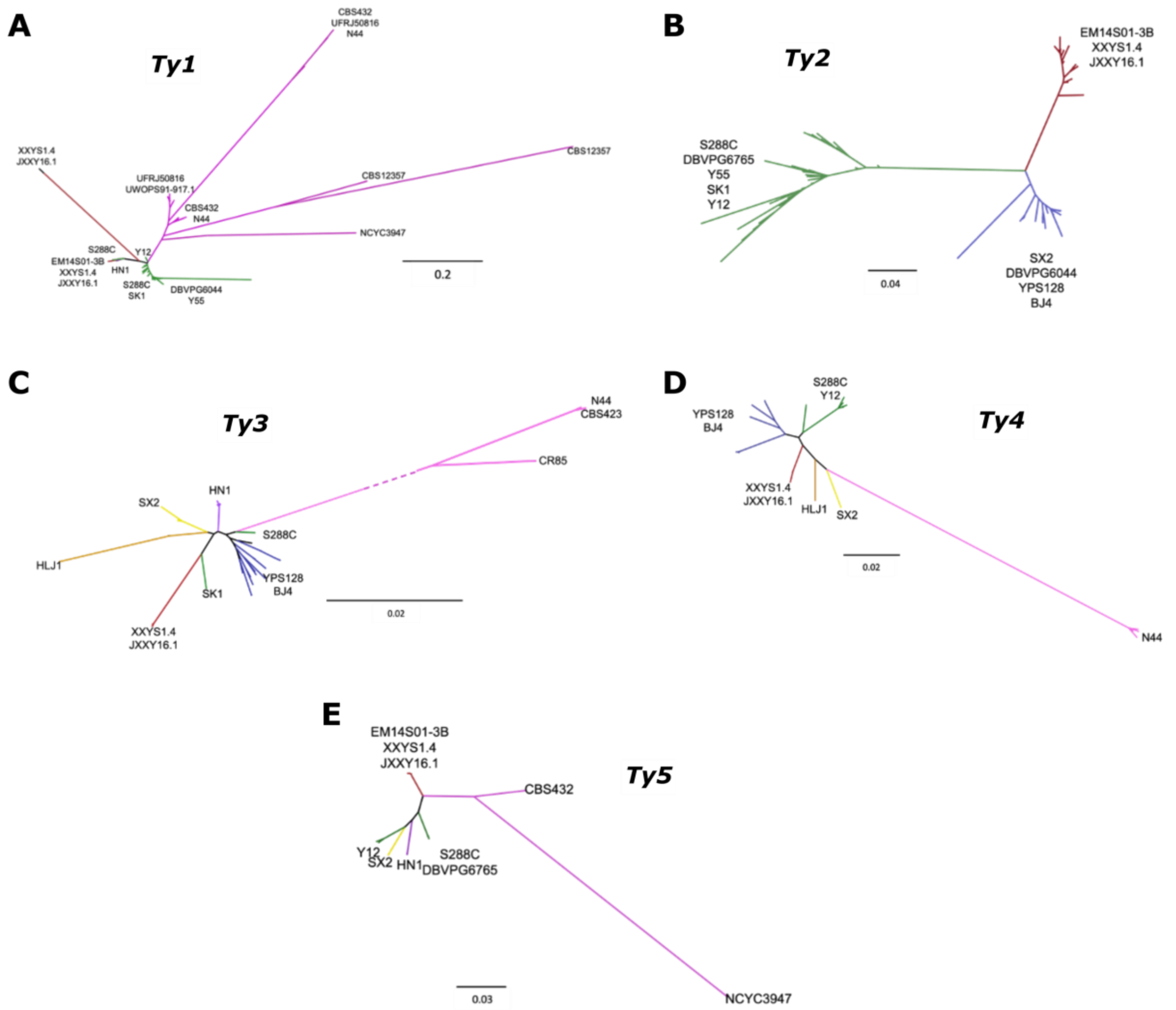


**Figure S21: Gene trees of *Ty* elements.** Maximum likelihood trees showing relationships between identified whole elements of **(A)** *Ty1*, **(B)** *Ty2*, **(C)** *Ty3*, **(D)** *Ty4* and **(E)** *Ty5*. Branches are colored according to rough groupings of strains with magenta indicating non-*S. cerevisiae* strains. Tips of branches are individual *Ty* elements labeled generally by the strain in which they were identified.

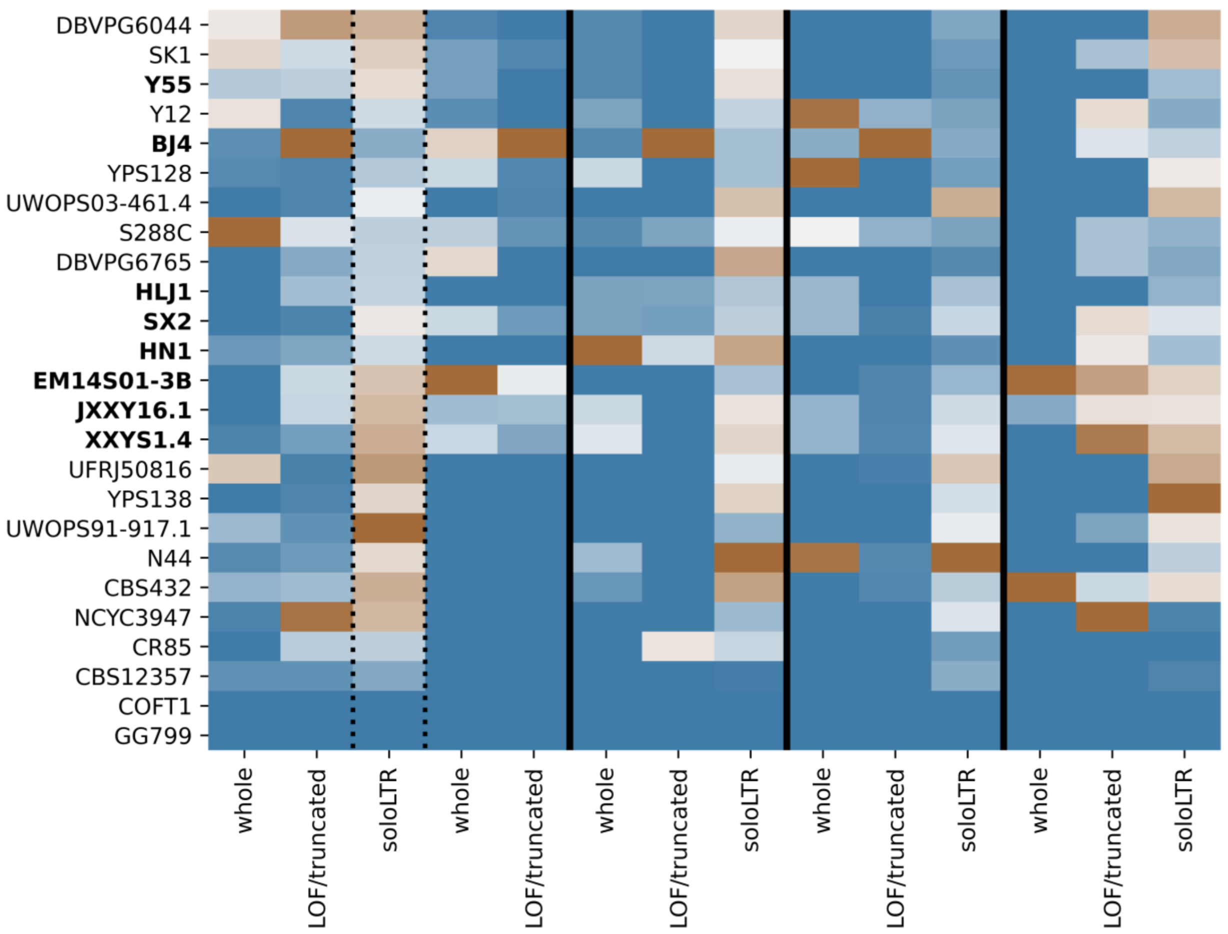


**Figure S22: Length of transposable elements in Saccharomyces** Total length of each type of transposable element found within each strain. Directly corresponds to the counts shown in **Fig. 3A**.

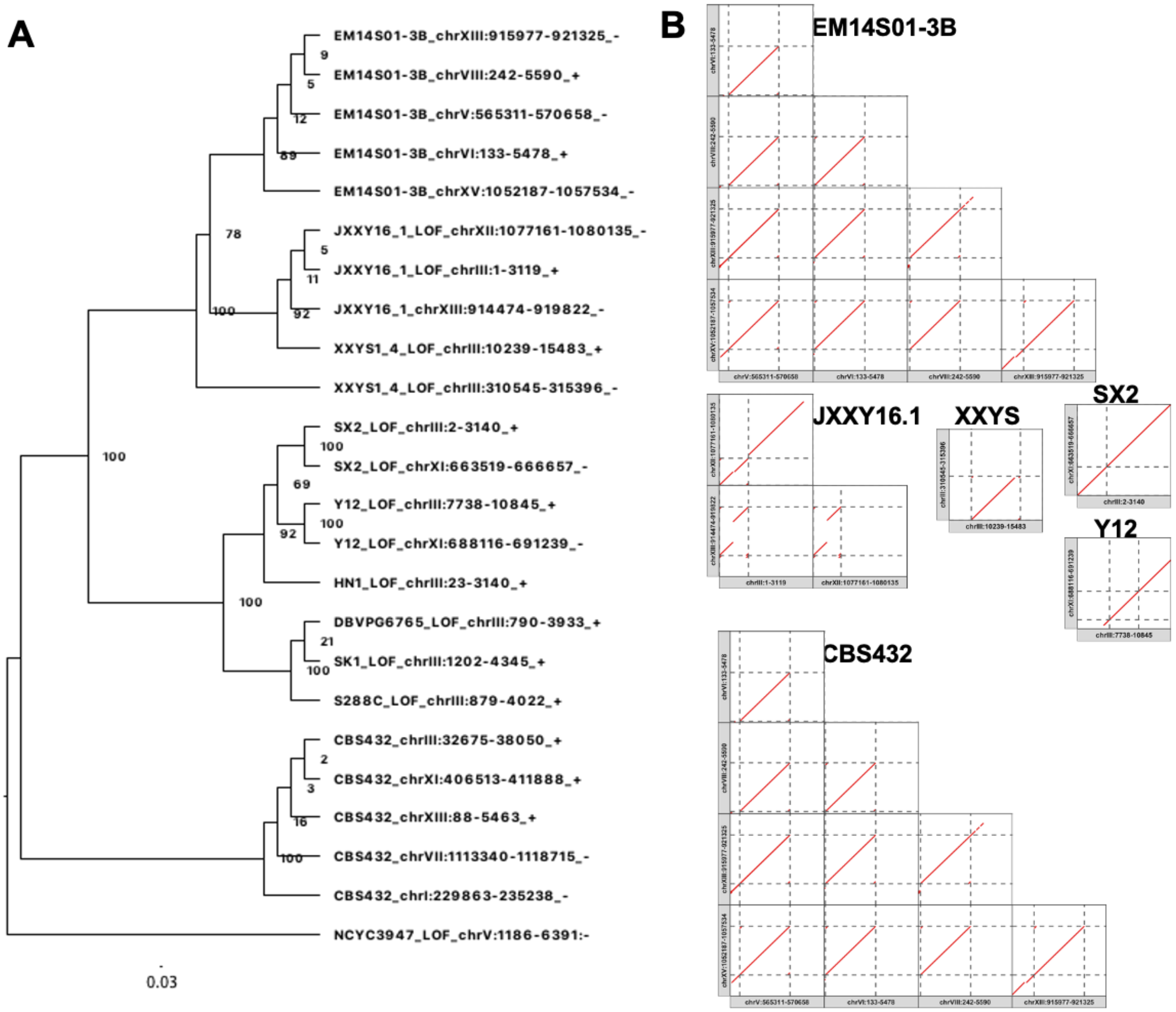


**Figure S23: *Ty5* element phylogeny and dotplots. (A)** Phylogenetic tree of *Ty5* elements. **(B)** Dot plots of *Ty5* elements. Dot plots shown alignments between =10kb regions within a strain containing different whole *Ty5* elements (dashed lines show the boundaries).

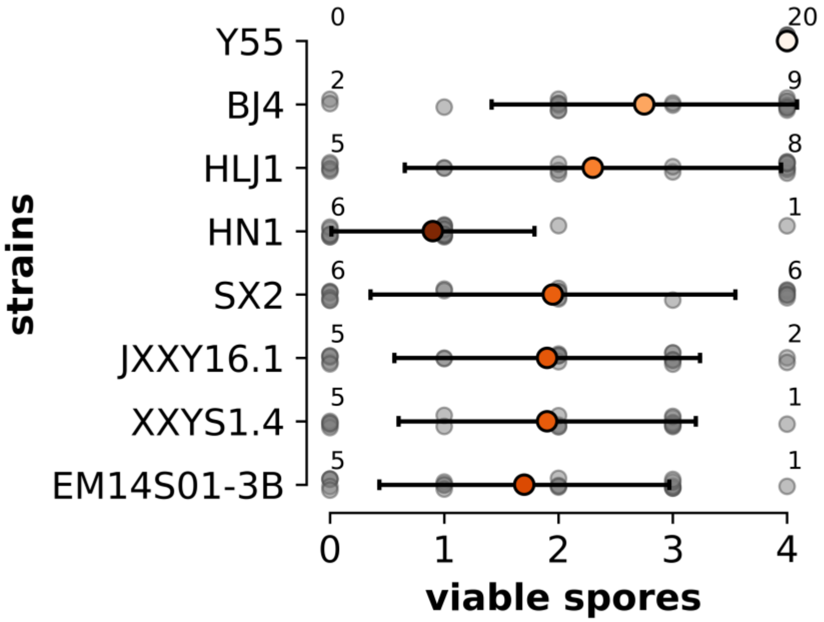


**Figure S24: Spore viability of intraspecific crosses.** The seven East Asian strains were mated with Y55, sporulated and the resulting tetrads dissected. For a positive control and reference Y55 was also mated with itself. 40 tetrads for cross were dissected and the number of viable spores is indicated by the grey dots. The total number of tetrads with 0 or 4 viable spores is indicated for each strain. Colored dots indicate the mean and standard deviation and are colored on a relative scale.
